# Using target capture to address conservation challenges: population-level tracking of a globally-traded herbal medicine

**DOI:** 10.1101/744318

**Authors:** Vincent Manzanilla, Irene Teixidor-Toneu, Gary J. Martin, Peter M. Hollingsworth, Hugo J. de Boer, Anneleen Kool

**Author notes:** Corresponding author: Vincent Manzanilla.

## Abstract

The promotion of responsible and sustainable trade in biological resources is widely proposed as one solution to mitigate currently high levels of global biodiversity loss. Various molecular identification methods have been proposed as appropriate tools for monitoring global supply chains of commercialized animals and plants. We demonstrate the efficacy of target capture genomic barcoding in identifying and establishing the geographic origin of samples traded as *Anacyclus pyrethrum*, a medicinal plant assessed as globally vulnerable in the IUCN Red List of Threatened Species. Samples collected from national and international supply chains were identified through target capture sequencing of 443 low-copy nuclear makers and compared to results derived from genome skimming of plastome, standard plastid barcoding regions and ITS. Both target capture and genome skimming provided approximately 3.4 million reads per sample, but target capture largely outperformed standard plant DNA barcodes and entire plastid genome sequences. Despite the difficulty of distinguishing among closely related species and infraspecific taxa of *Anacyclus* using conventional taxonomic methods, we succeeded in identifying 89 of 110 analysed samples to subspecies level without ambiguity through target capture. Of the remaining samples, we determined that eleven contained plant material from other genera and families and ten were unidentifiable regardless of the method used. Furthermore, we were able to discern the geographical origin of *Anacyclus* samples collected in Moroccan, Indian and Sri Lankan markets, differentiating between plant materials originally harvested from diverse populations in Algeria and Morocco. With a recent drop in the cost of analysing samples, target capture offers the potential to routinely identify commercialized plant species and determine their geographic origin. It promises to play an important role in monitoring and regulation of plant species in trade, supporting biodiversity conservation efforts, and in ensuring that plant products are unadulterated, contributing to consumer protection.

---

Human exploitation of biological resources is a major challenge for biodiversity conservation and sustainable development. Global trade and consumer demand for natural products provide increasing threats to species (Lenzen et al., 2012), and at the same time create markets in which regulation and authentication are extremely difficult (Newmaster, Grguric, Shanmughanandhan, Ramalingam, & Ragupathy, 2013; WHO, 2004).

Many traded plant products from wild populations are over-harvested. Increasing scarcity results in higher prices, and incentivises adulteration, substitution and poaching (Hamilton, 2004; Schippmann, Leaman, & Cunningham, 2002; Veldman et al., 2017). In recent decades, multilateral environmental agreements including the Convention on Biological Diversity (CBD) and the Convention on International Trade in Endangered Species (CITES) have addressed the trade of threatened species. In parallel, the World Health Organisation has developed guidelines for safety monitoring of herbal medicines in pharmacovigilance systems (WHO, 2004) and the Food and Agriculture Organisation, through the International Treaty on Plant Genetic Resources for Food and Agriculture (ITPGRFA), has set up a multilateral system to promote sustainable agriculture (Esquinas-Alcazar, 2004). However, implementing these regulations and guidelines is hampered by difficulties identifying plant products in trade. Multiple, complex and interacting supply chains can co-exist for a single plant product (A. Booker, Johnston, & Heinrich, 2012). Traded plant products are often not identifiable to species by their morphology or chemistry, as they may be dried, powdered, processed, or commercialised in mixtures with other ingredients. The design, implementation and enforcement of successful conservation actions as well as assessments of product authenticity and quality often require the identification of the geographic origin of species in trade. This is difficult as the development of efficient methods to identify and trace traded products are still in their infancy.

DNA barcoding has quickly gained popularity since Hebert et al. (2003) first advocated the use of short and variable DNA sequences, amplified using universal primers, for species identification and discovery of new taxa. DNA barcoding is highly effective for species-level identification in animals using a portion of the mitochondrial marker Cytochrome Oxidase 1 (COI) (P. D. Hebert, Hollingsworth, & Hajibabaei, 2016; P. D. Hebert, Ratnasingham, et al., 2016). In plants, standard DNA barcoding involves one to four plastid DNA regions (*rbcL, matK, trnH-psbA, trnL*), sometimes in combination with internal transcribed spacers of nuclear ribosomal DNA (nrDNA, ITS) (CBOL et al., 2009; Kress, 2017). Although these markers are very informative in many cases, no single marker or combination of markers routinely provide complete species-level resolution, especially in species-rich groups, let alone population-level assignment.

The development of high throughput sequencing (HTS) with new reagents and platforms expands the application of DNA barcoding in plants in a cost-effective fashion, in part because it removes the need of targeting short universal barcodes (Hollingsworth, Li, van der Bank, & Twyford, 2016; Lemmon & Lemmon, 2013). Two major approaches, shallow pass shotgun sequencing and target capture sequencing, have been proposed for increasing the coverage and resolution of plant DNA barcoding.

Shallow pass shotgun sequencing (commonly referred as genome skimming) is used to recover organellar genomes and nuclear ribosomal DNA sequences, increasing the amount of data per sample and leading to some increases in resolution (Manzanilla et al., 2018; Parks, Cronn, & Liston, 2009). Although workflows and bioinformatic pipelines are increasingly refined for this approach, current cost constraints mean that most genome skimming barcoding projects only have sufficient sequencing depth to generate comparative data for multi-copy regions such as plastid genomes and ribosomal DNA. These regions represent a limited number of independent loci, ultimately constraining resolving power (Soltis & Soltis, 2009; Wood et al., 2009).

Target capture sequencing offers the potential to overcome these deficiencies by efficiently targeting hundreds of low-copy nuclear markers, providing access to a much greater number of independent data points per unit of sequencing effort (Degnan & Rosenberg, 2009). Similar to genome skimming, target capture is successful in sequencing samples with poor DNA integrity (Brewer et al., 2019; Forrest et al., 2019) and allows sequencing hundreds of samples at the same time (Mamanova et al., 2010). It can also be designed to recover standard DNA barcodes in the same assay (Schmickl et al., 2016). Although target capture has been advocated as a powerful tool for molecular identification of plants (Pillon et al., 2013), its usefulness for determining and tracing the origin of plants in trade remained untested.

We evaluated the power of target-capture DNA barcoding by investigating the traceability of plant products reputedly derived from a vulnerable medicinal plant species, *Anacyclus pyrethrum* (L.) Lag. In addition to having a well-established international trade chain (Rankou, Ouhammou, Taleb, Manzanilla, & Martin, 2015), this species presents classic challenges for plant molecular identification such as recent radiation, frequent hybridization (Humphries, 1979) and large genome size (Garcia et al., 2013).

The genus *Anacyclus* (Asteraceae) comprises 12 species of annual and perennial weedy herbs with partly overlapping geographic ranges around the Mediterranean basin (Humphries, 1979; Rosato, Álvarez, Feliner, & Rosselló, 2017). Some species are abundant and have wide geographical ranges (for example, *A. clavatus* (Desf.) Pers. and *A. radiatus* Loisel.), whereas others are rare and have restricted ranges (for example, *A*. *maroccanus* (Ball) Ball and *A. pyrethrum* (L.) Lag.).

*A. pyrethrum* has a long history of use in traditional Arabic and Islamic, Ayurvedic and European medicine (Adams, Alther, Kessler, Kluge, & Hamburger, 2011; De Vos, 2010; Pittle, 2005). In the 13^th^ century, Ibn al-Baytār wrote that the plant was known across the world and traded from the Maghreb to all other areas (Leclerc, 1877). Its popularity as a medicinal plant stems from the many attested and putative pharmacological activities of its roots (Manouze et al., 2017). Traded from the Maghreb to India (Ved & Goraya, 2007) and Nepal (Tiwari et al., 2004), it is known to be over-harvested and is increasingly difficult to find in local markets in Morocco (Ouarghidi et al., 2013, 2012).

There are two accepted varieties, *A. pyrethrum* var. *pyrethrum* and var. *depressus*, both endemic to Algeria, Morocco, and southern Spain (Humphries, 1979; Rosato et al., 2017). In Morocco, *A. pyrethrum* var. *pyrethrum* is considered more potent and is up to ten times more expensive than var. *depressus* (Ouarghidi, Powell, Martin, de Boer, & Abbad, 2012). Both varieties are harvested from the wild and used extensively for the treatment of pain and inflammatory disorders across Morocco (Ouarghidi, Martin, Powell, Esser, & Abbad, 2013; Ouarghidi et al., 2012) and Algeria (Benarba, 2016; Ouelbani, Bensari, Mouas, & Khelifi, 2016), as well as the Middle East (Pittle, 2005) and the Indian sub-continent (Tiwari, Poudel, & Uprety, 2004). Although collectors are proficient in differentiating the two varieties, material traded as *A. pyrethrum* is adulterated and misidentified along the chain of commercialisation (de Boer, Ouarghidi, Martin, Abbad, & Kool, 2014; Ouarghidi et al., 2013, 2012).

We applied target-capture genomic barcoding to distinguish *Anacylus* species and geographical races in root samples in the national and international supply chains. We compare this novel approach with plastid genome and nrDNA ITS barcodes obtained from genome skimming data.

## Materials and Methods

### Sample collection

We made a reference collection of 72 accessions, consisting of 67 of *Anacyclus* and five of closely related genera, which includes 56 *Anacyclus* accessions and three of closely related genera we collected in Morocco and Spain during field work. We selected a total of 11 samples of *Anacyclus* and two of closely related genera from herbarium voucher specimens of species occurring elsewhere in the Mediterranean. Voucher specimen identifications, collection numbers and locality data are listed in Table S1 and the collection locations are mapped in Figure S1. We purchased 110 samples, each consisting of approximately 50g of roots from collectors, herbalists, middle-men, traditional healer, wholesalers, and export companies in Morocco and India (Table S2).

### Trade information

We obtained samples of commercialized species by asking collectors and traders in Morocco, for tiguendizt and iguendez, two of vernacular names used for both *A. pyrethrum* varieties (Ouarghidi et al., 2012). We asked traders in India for akarkara, a common name used there (Ved & Goraya, 2007). When acquiring the samples, we conducted semi-structured interviews about the roots’ trade with 39 informants, enquiring where the plant material was sourced, to whom it was sold and in what quantities, for what price, and if some types of roots were of higher quality than others. We followed the International Society of Ethnobiology Code of Ethics (“ISE Code of Ethics Online,” 2018), ensuring free, prior and informed consent, full disclosure and respect for the confidentiality of informants, during all interviews.

### Extraction and library preparation

We extracted DNA from reference and traded vouchers from approximately 40 mg of dry leaf or root material using the DNeasy Plant Mini Kit (Qiagen). Total DNA (0.2-1.0 μg) was sheared to 500 bp fragments using a Covaris S220 sonicator (Woburn, MA, USA) (Table S1-2). We prepared dual indexed libraries using the Meyer and Kircher protocol (Meyer & Kircher, 2010) for genome skimming and target capture (BioProject PRJNA631886).

### Target capture

Using the genome assembly of *A. radiatus* subsp. *radiatus*, we designed 872 low-copy nuclear markers and associated RNA probes by following the Hyb-Seq pipeline (Weitemier et al., 2014) (SI). For target capture enrichment, we prepared twelve equimolar pools with ten to 24 samples and an average 300 ng of input DNA per pool. The RNA probes were hybridized for 16 hours before target baiting, and 14 PCR cycles were carried out after enrichment following the MyBaits v3 manual. The enriched libraries and genome skimming libraries were sequenced on two Illumina HiSeq 3000 lanes (150bp paired-end).

### Data Processing

We retrieved four datasets from the genome skimming and target capture sequencing methods: (1) standard barcode markers (*matK*, *trnH*-*psbA*, the *trnL* intron and *rbcL*), (2) ITS, (3) complete plastid genomes (from shotgun genome skimming), and (4) hundreds of nuclear markers (from target capture) (Figure S2). The sequencing runs were trimmed and quality filtered using Trimmomatic (Bolger, Lohse, & Usadel, 2014). Low-copy nuclear markers and their alleles were retrieved for each sample. First, the reads were mapped against the selected low-copy nuclear markers (SI) using BWA v0.7.5a-r40 (Heng Li & Durbin, 2009). Duplicate reads were removed using Picard v2.10.4 (Wysoker, Tibbetts, & Fennell, 2015). Alleles were phased for each marker and individual using SAMtools v1.3.1 (H. Li et al., 2009). The last step of the pipeline combined the retrieved alleles into single gene matrices. We recovered plastome and ITS sequences by pooling shotgun and target enrichment sequencing data. Plastid genomes were built using MITOBim v1.8 (Hahn, Bachmann, & Chevreux, 2013). ITS sequences were recovered using BWA by mapping the reads to the reference ITS of *Anacyclus pyrethrum* (KY397478) for *Anacyclus* species and traded samples, to the reference ITS of *Achillea pyrenaica* Sibth. ex Godr. (AY603247) for *Otanthus* and *Achillea*, and to the reference ITS of *Matricaria aurea* (Loefl.) Sch.Bip. (KT954177) for *Matricaria* samples. During the mapping and the assembly steps, we retained sequences only with a minimum coverage of 10X.

### Phylogenomics

We aligned the recovered matrices (nuclear markers, ITS and plastomes) with MAFFT v7.471 (Katoh & Standley, 2013), refined with MUSCLE (Edgar, 2004) and filtered with Gblocks v0.91b (Talavera & Castresana, 2007). Phylogenies were inferred using RAxML v8.0.26 (Stamatakis, 2006), with 1000 bootstrap replicates under the GTRGAMMA model. For the low-copy nuclear markers, the species tree was inferred from the individual nuclear markers trees under the multi-species coalescence (MSC) framework with ASTRAL-III v5.5.9 (Zhang, Sayyari, & Mirarab, 2017). We used the multi-alleles option in ASTRAL-III for reconciliation of the independent evolutionary histories of the alleles. The molecular identification of traded roots was assessed from the MSC tree and posterior probabilities (PP) greater than 0.95. For the ITS phylogenetic reconstruction, we used additional Genbank references (Table S3).

We identified traded roots based on morphological characters (described in (SI)), and we used these to triangulate molecular identifications. Samples were categorised according to their position in the supply chain and geographical origin.

## Results

We constructed a reference database of DNA sequences from fresh and herbarium specimens, consisting of 83 individuals of 10 *Anacyclus* species, and 5 individuals representing outgroup species (Figure S1). We used this reference database to assess the identity and geographic origins of 110 root samples acquired from traded materials by comparing the results from the four different datasets (Figure 1). We show that the target capture approach is the most powerful method to identify plant species in trade and their geographic origin.

**Figure 1:**
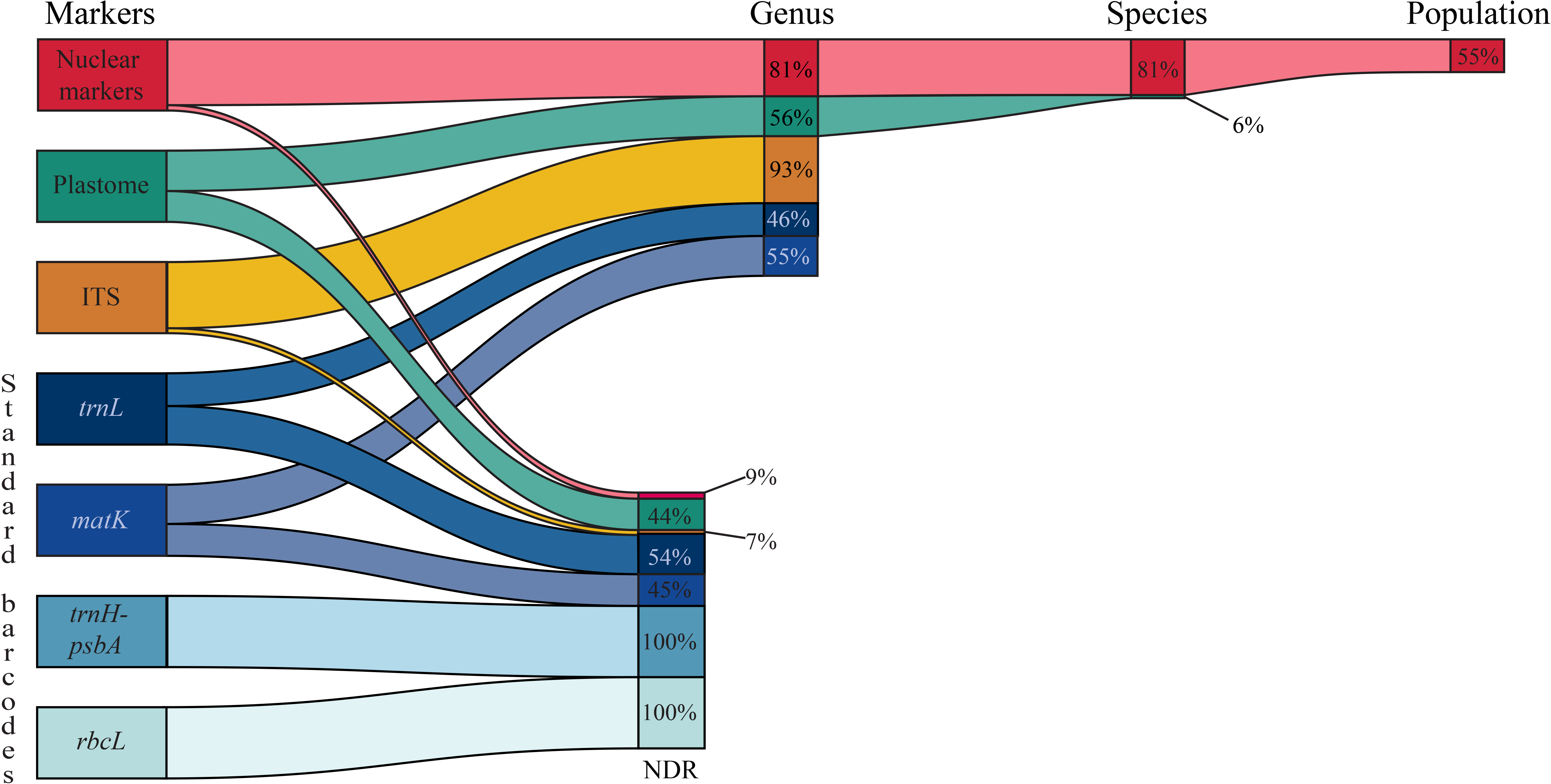
Sequencing recovery and identification success for the traded samples for each dataset. The figure shows the percentage of samples for which useful marker sequences were successfully retrieved for molecular identification for standard barcodes, ITS, plastomes and nuclear markers. No data recovered (NDR) is used to indicate samples for which no sequence data was recovered or where no identification at the genus level or below could be made. For the samples that produced useable data, the proportion of samples that resulted in identification at the genus, species and population levels is given.

### Data recovery for genome skimming and target capture data

After quality control filtering, an average 2.8 million reads (0.42 GB/sample) were obtained per sample from shotgun sequencing and 2.99 million reads (0.43 GB/sample) per sample for target capture (Figure S3, Table S4). The target capture yielded an average coverage of 303X for the 443 nuclear markers, whereas the unenriched genome skimming yielded an average coverage of 12X for the 443 nuclear data, 20X for the plastome data and 131X for the ribosomal data (Table S5). The loci coverage was calculated with *bedtools coverage* (v 2.29.2). Samples below an average of 50X coverage in the nuclear dataset, show a higher missing data rate (>7%) in the matrices. These samples were automatically discarded with our pipeline (Figure 2, samples in blue). Adulterated samples from other genera have a coverage close to zero (Figure 2, samples in orange, yellow and green). To obtain 100x coverage for the nuclear regions using a genome skimming approach, with an average genome size of the targeted species of 11.72Gb (Garnatje et al., 2011) and a duplication level of the genome skimming libraries of 9% (SI), it would require 14 HiSeq 3000/4000 lanes (Illumina, n.d.). From the genome skimming data we assembled ITS and the standard barcodes markers, as well as the plastome (Figure S2). Out of the 110 trade samples, we succeeded in assembling ITS for 102 samples (93%), the standard barcode regions for 51 to 61 samples (46% to 55%), and plastomes for 49 samples (44%) (Figure 1). The resulting aligned matrices for each of the datasets were 633 bp for ITS (including 5.8S), 4408 bp for the standard barcoding regions, 110,003 bp for the plastome and 289,236 bp for the 443 nuclear markers recovered from the target capture approach (Table S1). The standard barcoding regions included the full coding regions of *matK* 1523 bp and *rbcL* 1438 bp, as well as *trnH-psbA* 500 bp and *trnL* 947 bp. The bioinformatics workflow for data analyses is described in Figure S2.

**Figure 2:**
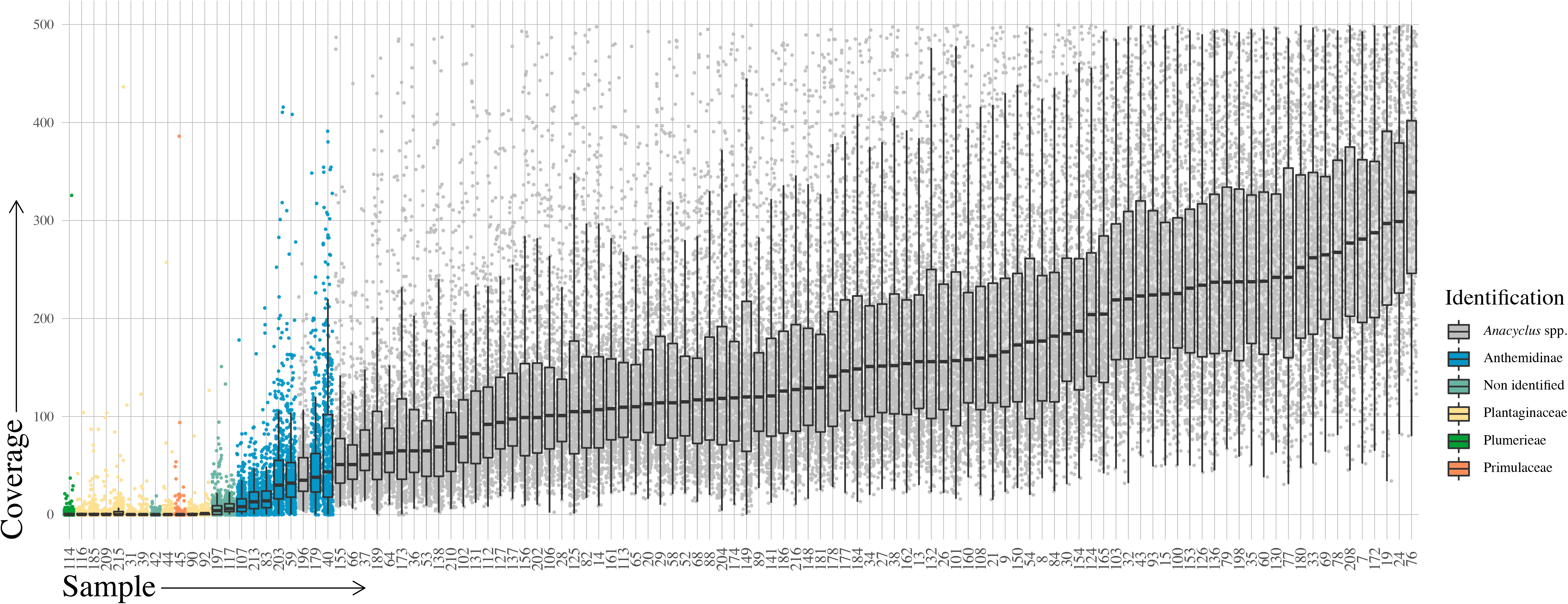
Box-and-whisker plots showing retrieved coverage of 443 targeted markers for each sample. Identified and non-identified samples are colour coded.

### Comparative levels of species discrimination using different approaches

The ITS, plastome, and standard barcodes phylogenies highlight the complex evolutionary history of *Anacyclus*. The ITS phylogeny lacks resolution in general (Figure S4-5). The outgroups *Tanacetum*, *Matricaria*, *Achillea*, *Othanthus* and *Tripleurospermum* have well-supported bootstrap values, but within the genus *Anacyclus*, only *A. atlanticus* Litard. & Maire, *A. maroccanus* and *A. radiatus* are highly supported. The plastome phylogeny shows very good support at genus level for the *Anacyclus* node, and at species level for the outgroups. The lack of variation in the plastid genome within the genus *Anacyclus* results in little phylogenetic support with no species-specific clusters recovered (Figure S6-7). The standard barcode regions, *matK*, *rbcL, trnH*-*psbA* and *trnL* (Figure S8-11) displayed low levels of resolution at the species level, even using the full coding regions of *matK* and *rbcL* (e.g. rather than the 800-900 bp of *matK* and 654 bp of *rbcL* typically recovered using standard barcoding primers;(Alsos et al., 2020; Hollingsworth, Graham, & Little, 2011)).

The 443 nuclear markers recovered by target capture, led to a well-resolved phylogeny and high levels of species discrimination: all the genera in the Matricariinae tribe and all interspecific relationships are well-supported, with most nodes showing posterior probabilities (PP) of 1 (Figure 3 and S12. Within *Anacyclus*, all species, sub-species and varieties are well supported. PP are lower for *A. monanthos* (PP = 0.75). The complex of hybrid species composed of *A. clavatus*, *A. homogamos*, and *A. valentinus* is polyphyletic and shows signs of hybridization and incomplete lineage sorting. Intraspecific nodes have PP varying between 0.27 and 1, mostly depending on species population structure.

**Figure 3:**
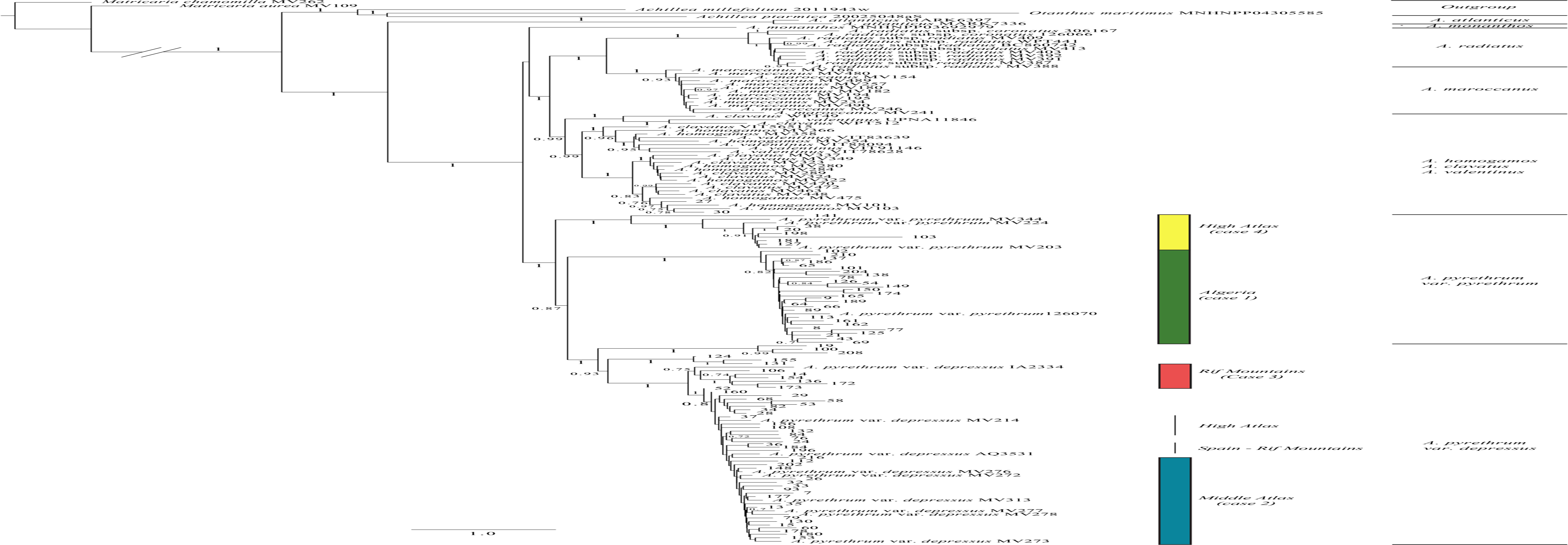
Multispecies coalescent phylogenetic tree of the nuclear loci dataset. The reference dataset includes taxon labels and herbarium accession numbers and the traded samples are numbered. Supported clades with an associated geographic origin are indicated by coloured bars.

### Assessment of *Anacyclus* trade

Interviews with 39 harvesters, middlemen, retailers, and wholesalers in various Moroccan cities, indicate that the national and international trade of *Anacyclus pyrethrum* follow two separate supply chains (SI). Retailer herbalists in Moroccan cities are supplied by middlemen who acquire the plant from local harvesters from rural communities. These retailers typically hold between a few hundred grams to one kilogram of the plant material in their shops. In contrast, wholesalers who export the plant internationally, hire professional harvesters who travel across the geographical range of the species to collect plant material. Harvested roots are brought directly from the wild to the export companies in Rabat, Casablanca and Tangier, from where they enter the international market, including supply of material to India. According to informants from export companies, between 3-10 tons of the plant product can be stocked at a time.

Our examination of material in trade involved screening a total of 66 bags each containing an average of 25g of dry roots. Initial morphological examination of these samples identified obvious non-*Anacyclus* adulterants in 39/66 batches. The adulterants were present with a proportion from 3% to 100% with an average of 42%. The non-*Anacyclus* adulterants were found at high frequency in collections from traditional healers and herbalists, less so from collectors, wholesalers and export companies (Figures 4–5, Table S6).

**Figure 4:**
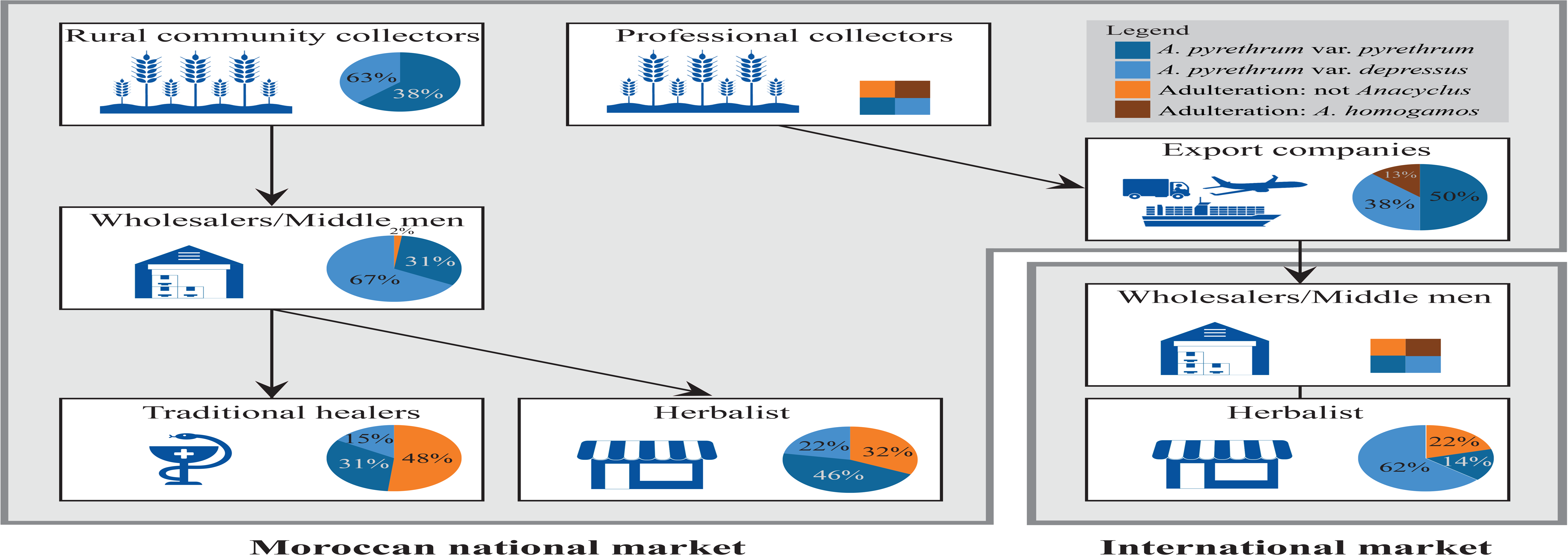
National and international supply chains of *A*. *pyrethrum*. Pie charts represent the proportion of *A. pyrethrum* (light and dark blue represent var. *depressus* and var. *pyrethrum* respectively) and adulterated samples (orange and brown for *A. homogamos* and other adulterants) by each stakeholder. We were unable to obtain samples from wholesalers/middlemen in India or professional collectors in Morocco (indicated by square boxes).

**Figure 5:**
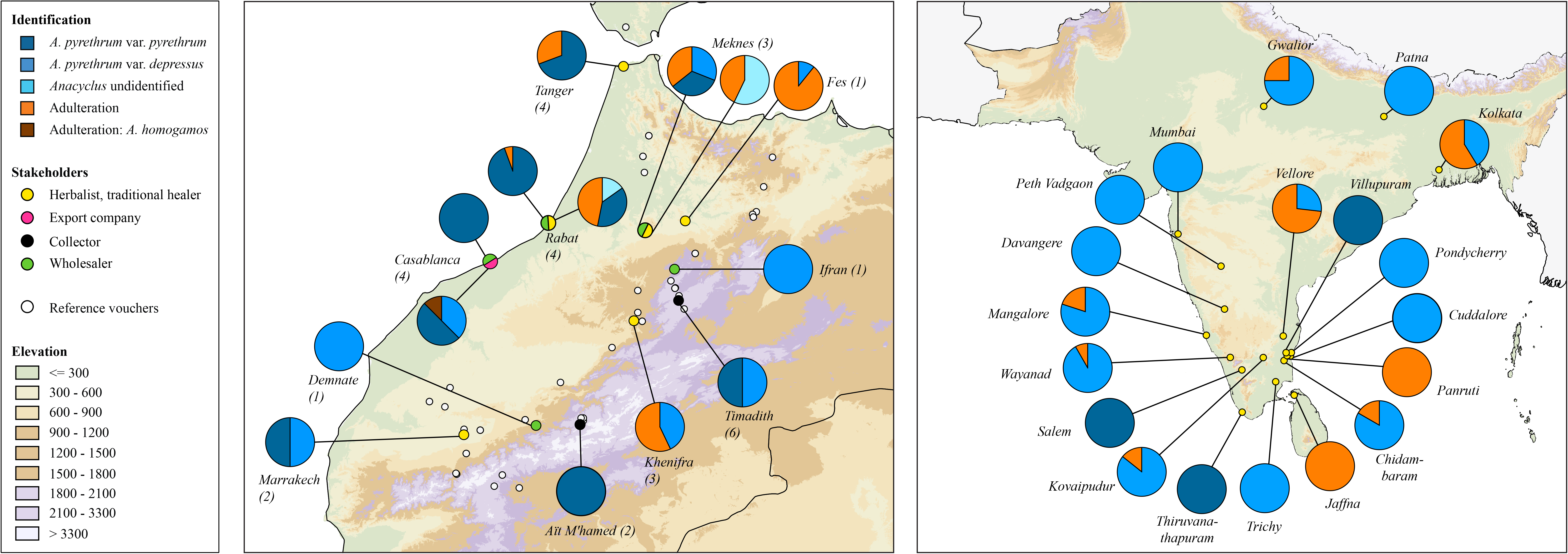
Species identification of market samples. Sample locations are shown with coloured circles according to the type of stakeholder. A pie-chart with the proportions of adulteration and identified species is represented for each location in (a) Morocco (native range) and (b) India (exported material).

We selected 110 individual roots for DNA analysis from the 66 root batches. Of these 99 had a morphology consistent with *Anacyclus*, and 11 which were classed as similar to *Anacyclus* but likely to be non-*Anacyclus* based on their morphology. We recovered partial plastome assemblies of these 11 non-*Anacylus* roots and using sequence queries against GenBank, we obtained identifications for nine *Plantago* spp., *one Primula* spp. and one *Plumeria* spp. (Figure S13, Table S2).

Of the 99 *Anacyclus* roots, 10 had no identifiable DNA sequences via any of our methods, with 89/89 of the remaining samples identified through target capture (Figure S13). The ten discarded samples presented very fragmented plastome assemblies with average 89% missing data in the nuclear matrices. These samples had a very low DNA integrity.

The plastome sequences enabled identification of seven roots to the species level, with the remainder identified as *Anacyclus* sp. (Figure 1, Table S7). Neither ITS nor any of the standard barcode regions were able to discriminate any of these samples below the genus level (Figure 1, S4-5, S8-11).

The nuclear markers gave much higher resolution within *Anacyclus* (Table S2, S7). In our investigation of plant material traded within Morocco, our analyses of six individual root samples from four rural community collectors identified three *Anacyclus* var. *pyrethrum* and three var. *depressus*. Our analysis of five samples from three wholesaler ‘middle-men’ in Morocco identified two *Anacyclus* var. *pyrethrum* and three var. *depressus*. Our sequences from 19 samples from 10 herbalists revealed 12 *A. pyrethrum* var. *pyrethrum* and seven var. *depressus*. Likewise, our 11 samples from six traditional healer sources identified seven *A. pyrethrum* var. *pyrethrum*, and four var. *depressus*. For material traded in international markets, our 17 sequenced samples from three export companies in Morocco identified nine *A. pyrethrum* var. *pyrethrum*, six var. *depressus*, and two *A. homogamous*. Our analysis of 30 samples from 17 herbalists in India identified three *A. pyrethrum* var. *pyrethrum* and 27 var. *depressus*.

### Geographical source

Market samples in Morocco originate both from various Moroccan areas as well as Algeria, and material from all of these populations of origin can be found in Indian market samples (Table S1, Figure 3, 6). Of the 99 non-adulterated roots which we identified to species level using target capture, we were able to associate 67% to a specific geographic region (Figure 1, 3, Table S1). Using phylogenetic analysis, root samples of *A. pyrethrum* var. *pyrethrum* clustered with reference material from the High Atlas (Figure 6, case 4), and *A. pyrethrum* var. *depressus* roots clustered with reference material from different regions in Morocco, including the Rif Mountains, the High Atlas and the Middle Atlas (Figure 6, case 2 and 3). Evidence for the international trade from Algeria to Morocco is highlighted by a distinctive clade that includes traded roots collected from west Algeria (Figure 6, case 1). The geographic origin was only resolvable with target capture data; standard barcoding regions, ITS and plastome data lacked variation, resolution or both (Figure 1).

**Figure 6:**
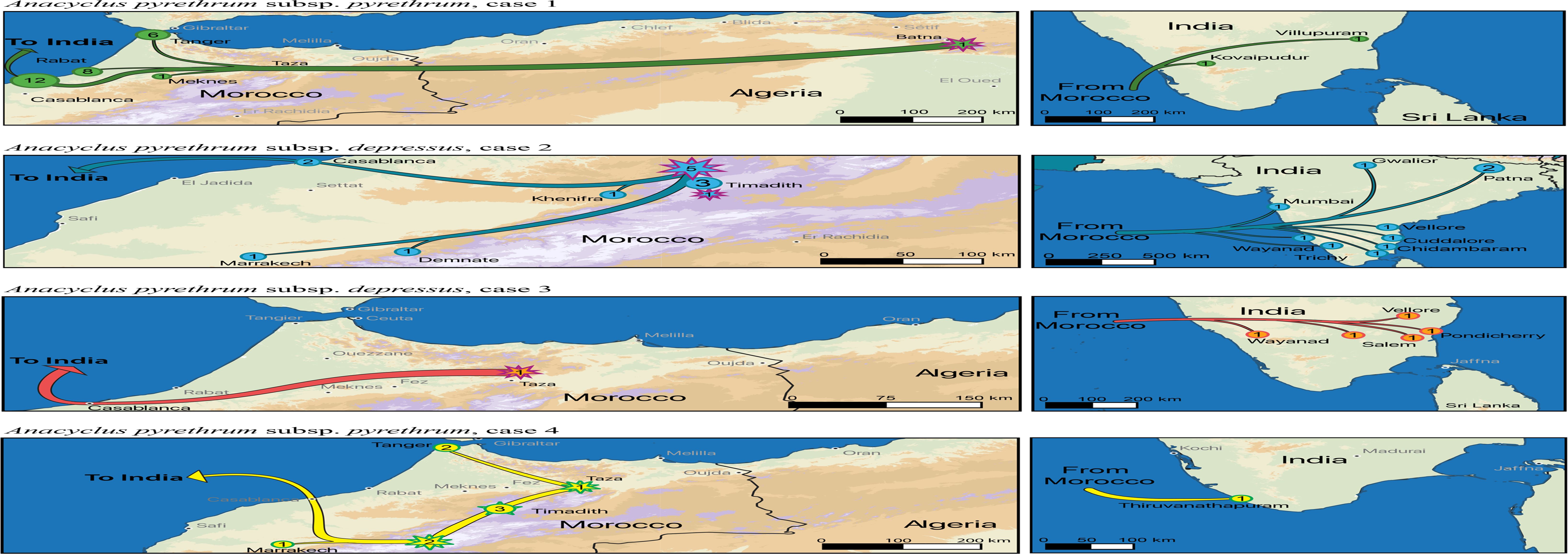
Map of the origin of the traded samples. The figure relates four supported clades mentioned in the Figure 3. The stars indicate reference samples and the dots the traded samples. The number in the circle or the start corresponds to the number of individuals.

## Discussion

This study illustrates the potential for target-capture based DNA barcoding to form the next wave of standard plant DNA-barcoding tools and provide the greatly needed species-level resolution. A key rate limiting step for the standard plant barcodes is that they are fundamentally recovering data from just one or two independent loci (plastid DNA and ITS), which often show trans-specific polymorphism and barcode sharing among related species (Hollingsworth, Graham, & Little, 2011). Even using complete plastid genome sequences suffers from the same problem, as the data are all physically linked in a single non-recombinant uni-parentally inherited locus. Several recent hybridization events have occurred in the *Anacyclus* genus and entire plastid genomes or plastid barcode markers, and/or ITS provide limited resolution below genus level (Figure 1). In contrast, our target capture approach using hundreds of nuclear markers yields significantly higher molecular identification success and more accurate resolution to species and even population level (Figure 1).

### *Species identification, species in trade, and geographic origins of* Anacyclus

These data provide new insights into trade of *A. pyrethrum* and highlight the extent of adulteration and the scarcity of *A. pyrethrum* var. *pyrethrum* (Figures 2–3, 6). Only a small proportion of the tested samples from herbalists and traditional healers were the potent *A. pyrethrum* var. *pyrethrum*, with the Indian market in particular dominated by var. *depressus.* In both Morocco and India some individual sellers had entirely or almost entirely adulterated products. Most of the non-*Anacyclus* roots we sequenced were identified as *Plantago* spp. by shotgun sequencing, despite being sampled from six different localities including Morocco and India. As *Plantago* roots are similar in appearance to *Anacyclus*, it is possible that they are deliberately added as a ‘difficult to identify’ adulterant which may go unnoticed by non-specialists.

Our analysis of samples from collectors, wholesalers and export companies detected much less adulteration at this point in the supply chain (Figure 4, Table S6). Collection of *A. pyrethrum* var. *pyrethrum* is carried out by professional harvesters employed by export companies who travel across the country and are considered poachers by local communities (Ouarghidi, Powell, Martin, & Abbad, 2017). Local harvesters have increasing difficulty to supply local trade chains (Ouarghidi et al., 2017, 2012), which may finally result in increased adulteration rates in the poorly-governed, national value chains (Figure 4), as has also been observed elsewhere (A. Booker et al., 2012). Our results also identify previously unreported international trade in North Africa prior to export to the Indian sub-continent (Figure 3, 6). We provide evidence that export companies in Morocco source material not only in this country, but also from neighbouring Algeria (Figure 6). Applying this molecular identification approach enables us to distinguish samples at population level and uncover these hidden international sourcing channels.

#### Conservation of *A. pyrethrum*

High national and international demand for *A. pyrethrum* likely encourages its overharvesting and adulteration. As the plant is a remedy of the Indian pharmacopoeia, its demand is likely to increase along with that of other Ayurvedic medicines (Kala, Dhyani, & Sajwan, 2006). Although *A. pyrethrum* has been assessed to be vulnerable internationally (Rankou et al., 2015) and endangered in Morocco on the IUCN Red List, the plant is not listed in the CITES appendices and its international trade is not regulated. Nonetheless, continued overharvesting is driving wild populations to critical levels and conservation policies are necessary. Common strategies to conserve overharvested medicinal plants often include collection and trade restrictions as well as cultivation (Schippmann et al., 2002). Cultivation is often proposed as a solution to both conservation issues and sourcing high quality, appropriately identified material (Hamilton, 2004; Schippmann, Leaman, & Cunningham, 2006; Schippmann et al., 2002). The cultivation necessitate engagement with local communities that depend on plant harvest, as well as monitoring of professional trading networks. Both promotion of sustainable harvesting for livelihood security as well as restriction of unsustainable professional trade are needed. Only with fine-grained mapping of sourcing areas and supply chains, as our results highlight for *A. pyrethrum*, can meaningful conservation action be achieved. With the implementation of target enrichment, we identified the origin of the harvested populations of medicinal or traded populations and point out where conservation efforts should be firstly implemented. We also reveal a previously undocumented harvesting and trade in Algeria (Figure 6, case 1), and show that *Anacyclus* is harvested at a national scale in Morocco (Figure 6, case 2-4). This study gives scientific evidence to support conservation programs like GDF’s High Atlas Cultural Landscapes Programme and potentially attract more attention for future conservation projects for *Anacyclus*.

### Future prospects for plant DNA barcoding

Key criteria for developing new DNA barcoding approaches include resolving power (telling species apart), recoverability (enabling use on a wide diversity of tissue sources), and cost and efficiency (enabling scaling over very large sample sets).

In terms of resolving power, the target capture approach used here offers substantial improvement compared to plant barcodes based on plastid sequences and ITS. The key enabling step is access to multiple nuclear markers, as this reduces sensitivity of the identification to introgression of one or two loci as is the case for barcodes from rDNA or the plastid genome. In this study, we show high resolution from nuclear markers retrieved by target capture for identification of species in a genus that has undergone recent hybridization events (Manzanilla, 2018). The successful recovery of these markers from low quality input DNA is also important and combined these observations make the case that this method provides a viable solution for future plant barcoding. In contrast, shotgun sequencing requires a substantial sequencing effort to retrieve the same suite of nuclear markers for species with large genomes. Thus for *Anacyclus* it would require 14 HiSeq 3000 lanes to obtain the same coverage of the loci we have used here. Overall, and regardless of approach, our findings provide empirical evidence to support predictions made in previous review papers (Hollingsworth et al., 2016) that barcoding based on nuclear markers should outperform standard barcoding methods.

The successful recovery of sequences via target capture depends on how closely related the sampled species are to the reference set on which the baits were designed, and/or the level of variation in the loci that form the bait set (McCormack, Tsai, & Faircloth, 2016; Paijmans, Fickel, Courtiol, Hofreiter, & Förster, 2016). In the current study, the baits were designed from *Anacyclus* and related genera. This optimized their specificity to our study group and enabled their successful high-resolution application for assessing trade. A clear challenge for wider use of target capture, is applicability over much greater phylogenetic distances. The recently published universal angiosperm baits (Buddenhagen et al., 2016; Johnson et al., 2018), designed to recover 353 loci from a wide diversity of flowering plants offer great potential here. The angiosperm universal bait kit (Johnson et al., 2018) can provide species and population-level resolution (Van Andel et al., 2019). In addition, there is a need for a more general evaluation of when, how and at what scale to most effectively combine taxon-specific bait sets (as used here) with universal bait sets, to simultaneously obtain very high resolution and sequencing success over wide phylogenetic distances.

Another important aspect of recoverability is efficacy with degraded DNA. Drying, storage, and transportation affect the quality of plant material in trade and can cause extensive DNA degradation (Anthony Booker et al., 2014). In consequence, traded samples have similar challenges to working with ancient DNA, herbarium samples or archaeological remains. Target capture is particularly well suited to this challenge (McCormack et al., 2016; Paijmans et al., 2016). Although shotgun sequencing can also be very effective on degraded material (Alsos et al., 2020; Bakker, 2017; Zeng et al., 2018), our recovery rate in this study was greater for target capture than shotgun sequencing (Figure 1), and we recovered data from 100% of the samples used for establishing the reference library and over 70% of the samples in trade. This 70% success rate for hundreds of nuclear markers providing high resolution from suboptimal tissue of traded samples is noteworthy. The other mainstream approach for highly degraded DNA is a portion of the chloroplast *trnL* (UAA) intron, specifically the P6 loop (10–143 bp)(Taberlet et al., 2007). This has been highly successful in recovering sequence data from degraded samples (Parducci et al., 2012; Willerslev et al., 2014). However, this short region of the plastid genome has a low variation at the species level and does not typically discriminate among con-generic species (Taberlet et al., 2007).

The target capture methodology presented here, including library construction and sequencing, cost 70 USD per sample in 2016, a price similar to those presented by Hale et al. (2020). With the optimisations suggested in Hale et al. (2020), prices drop to 22 USD per sample. Thus, this target enrichment approach is quickly becoming affordable for large-scale biomonitoring projects. With a growing interest and investment in DNA-based identification solutions in medicine and industry (Afshinnekoo et al., 2017; Menegon et al., 2017), ongoing work is expected to continue to optimize protocols and drive these costs down even further, reaching those of standard barcoding approaches (Hebert et al., 2018).

## Conclusion

In plants, the frequent sharing of plastid and ribosomal sequences among con-generic species, coupled with the difficulty of routinely accessing multiple nuclear markers, has acted as a constraint on the resolution of DNA barcoding approaches. Current advances in sequencing technology and bioinformatics are removing this constraint and offer the potential for a new wave of high-resolution identification tools for plants. These approaches, such as target capture have the capability to distinguish species and populations, providing insights into diversity and ecology, as well as the multitude of societal applications which require information on the identification and provenance of biological materials.

## Supporting information

SI Manzanilla et al. 2021

## DECLARATIONS

### Authors’ contributions

The project was coordinated by AK, GM, HdB and VM. VM did the design of the study and performed data analysis. AK, HdB, ITT, PH and VM wrote the manuscript. All authors provided useful contributions to data analysis and interpretation of the results. All authors have read and approved the final version of the manuscript.

## Acknowledgments

Maxime Borry for the DNA extraction; Julie Hawkins for hosting V.M. in Reading during the phylogenomics analyses; the Global Diversity Foundation team in Morocco for their logistic support during the field work. The PET group for their useful discussions and recommendations during the project. This work was performed on the Abel Cluster, owned by the University of Oslo and the Norwegian metacenter for High Performance Computing (NOTUR), and operated by the Department for Research Computing at USIT, the University of Oslo IT-department, http://www.hpc.uio.no/.

## Reproducibility

For reproducibility purposes, all the scripts used during the data processing are available on the OSF work repository https://osf.io/9bh3p/. New sequencing data have been deposited under a single NCBI BioProject accession PRJNA631886.

