## Supplementary material for "Using target capture to address conservation challenges: population-level tracking of a globally-traded herbal medicine": SI Manzanilla et al. 2021

Corresponding author: Vincent Manzanilla

### SI Material and Methods

**Morphological estimation of adulteration.** To estimate adulteration, collected trade samples were screened for plant parts that could be morphologically identified. Identifications were supported by data from previous studies on adulterants for *A. pyrethrum* (de Boer, Ouarghidi, Martin, Abbad, & Kool, 2014; Kool et al., 2012). Based on the shape, colour, and morphological structure of the root cross-section, roots were grouped into 'possibly belonging to the genus *Anacyclus*' or 'definitely adulterated', and the two subsets were subsequently weighed. All roots identified as 'possibly belonging to the genus *Anacyclus*' species were selected for molecular identification.

**Marker design, skimming data and denovo assembly.** Nuclear markers for molecular identification of the Matricariinae sub-tribe (Asteraceae) were designed using novel skimming data of an accession of *A. radiatus* subsp. *radiatus* (voucher MV54) (Table S1). The Hyb-Seq pipeline (Schmickl et al., 2016) was used to find conserved DNA regions of sufficient length and to exclude multiple copy genes and transposable elements. The *A. radiatus* subsp. *radiatus* voucher (MV54) yielded 2.86 µg of DNA measured on a Qubit 2.0 fluorometer (Invitrogen/Life Technologies, Carlsbad, CA, USA). DNA integrity and concentration were assessed using a Fragment Analyser (Advanced Analytical, Heidelberg, Germany) and the High Sensitivity genomic DNA Reagent Kit (50–40,000 bp). The sample was sequenced on an Illumina NextSeq 500 paired-end system using a TruSeq DNA PCR-Free library kit. Library adapter sequences and low quality reads were removed with Trimmomatic v. 0.32 (Bolger, Lohse et al. 2014) with a quality threshold set at Q20 with a sliding window of 10 bp.

Prior to the *denovo* nuclear genome assembly, the plastid genome was assembled with the trimmed reads using MITObim v1.8 (Hahn, Bachmann, & Chevreux, 2013) using the plastid genome of *Chrysanthemum indicum* L. (NC_020320) as a reference. Protein-coding genes in the plastid genome were annotated with DOGMA (Wyman, Jansen, & Boore, 2004), and after visual inspection, a gene map was drawn using OGDRAW v1.2 (Lohse, Drechsel, & Bock, 2007). Selective filtering of the plastid and mitochondrial genomes as well as nrDNA were done using BWA v0.7.5a (Langmead & Salzberg, 2012). The previously assembled plastid and mitochondrial genomes of *Helianthus annuus* L. (NC_023337.1) and the nrDNA from *Anacyclus valentinus* L. (GU818490) were used as references. The nuclear genome of *A. radiatus* subsp. *radiatus* was assembled using SOAPdenovo2 vr223 (Xie et al., 2014) with nine kmer values between 20 and 100. The best genome assembly was determined using Quast v2.3 (Gurevich, Saveliev, Vyahhi, & Tesler, 2013).

Low-copy nuclear markers (600-1000 bp in length) were identified using the Hyb-Seq pipeline based on the skimming assembly of *A. radiatus*, and the transcriptome assembly of a close relative outgroup, *Matricaria matricarioides* (Less.) Porter (voucher ALTA132745) (Matasci et al., 2014). The original Hyb-Seq pipeline was adapted to identify introns as well as exons (Schmickl et al., 2016). The modified script used for the selection of the low-copy nuclear markers is available on Open Science Framework, *Anacyclus* project folder (https://osf.io/9bh3p/).

Transcriptome and skimming data were pre-processed to ensure selection of sufficiently long nuclear regions as markers using the Hyb-Seq pipeline, filtering out plastid mitochondrial sequences using *Helianthus annuus* NC_023337.1 as a reference, and nrDNA with the reference of the *A. radiatus* assembly. A length threshold was applied on the remaining data and transcripts below 120 bp (RNA probe size) and contigs from the skimming data below 600 bp were discarded. Subsequently, the contigs were mapped against the *M. matricarioides* transcriptomes using Blat v3.5 (Kent, 2002), and alignments were selected with a minimum length of 80% of the contig size. Alignments with more than 10% divergence and contigs with more than one match against the *M. matricarioides* transcriptomes were discarded. The obtained preliminary set of markers was mapped with the Burrows-Wheeler Aligner (BWA) v0.7.5a-r405 (Li & Durbin, 2010) against the reads from the *A. radiatus* nuclear genome assembly. We extracted the coverage from this alignment using BEDtools v2.17 (Quinlan, 2014), and contigs with a higher coverage than average were discarded because they were suspected to be multiple copy genes or contain transposable elements. A total of 872 putative low copy nuclear markers were retained. To ensure that the probes targeted only the nuclear genome, we mapped the probes against the previously assembled organelles and nrDNA with BWA and discarded those that matched. From the selected loci, 19,246 RNA probes with a length of 120 bp and a tilling density of 3.4X were designed and produced by Arbor Bioscience (Ann Arbor, Michigan, USA).

**Nuclear genes filtering.** Nuclear gene trees were reconstructed for each individual nuclear locus. Samples with >7% missing data across markers were removed from the entire dataset, as well as markers with >5% missing data, as these were considered to have insufficient enrichment success. Retained matrices were re-aligned using MUSCLE v3.8.31 (Edgar, 2004) and filtered with Gblock (Talavera & Castresana, 2007). The final set consisted of 443 low-copy nuclear markers with two alleles per individual, with a minimum length of 400bp, no missing samples and less than 5% missing data. For each of these markers, we inferred a gene tree using RaxML v8.0.26 (Stamatakis, 2006) with 1000 bootstrap replicates under the GTRGAMMA model. Out of the 872 nuclear markers, 429 were discarded because we suspected to have transposons within these markers. Plants with huge genomes are known to have many transposon and low-complexity regions. Without a fully annotated genome, like in this case, it is difficult to assess quality of the designed markers before sequencing them.

**Results.** The best assembly of the *A. radiatus* subsp. *radiatus* genome was obtained with kmer values set at 70 bp, resulting in an N50 score of 869 bp with a total contig length of 1.4 Gbp. Assuming that the genome size of *Anacyclus* is 16 Gb (Bennett & Leitch, 2005; Humphries, 1981), the assembly represents approximately 8% of the *A. radiatus* subsp. *radiatus* genome. Raw data are deposited in NCBI Bioproject PRJNA631886. The plastid genome assembly yielded a final annotated genome of 150,925 bp with an average coverage of 350X (Table S1).

### SI Figure and table captions

**Fig. S1**. Geographical mapping of the sampled populations. Dots show the locations of the different populations and the colors represent different species.

**Fig. S2**. Bioinformatics workflow to retrieve the ITS, plastome, standard and nuclear marker datasets.

**Fig. S3**. Number of reads per sample before and after trimming.

**Fig S4**. Maximum Likelihood ITS phylogenetic tree reconstructed from the reference dataset only.

**Fig. S5**. Maximum Likelihood ITS phylogenetic tree with reference and traded samples. GenBank reference sequences are written in plain text, trade samples are in grey and reference samples from our study in bold. The vertical lines on the right of the phylogeny represent supported clades with the associated species names.

**Fig. S6**. Maximum Likelihood plastome phylogenetic tree reconstructed from the reference dataset only.

**Fig. S7**. Maximum Likelihood plastome phylogenetic tree with reference and traded samples. Traded samples are indicated in the phylogram by their accession number.

**Fig. S8**. Maximum Likelihood matK phylogenetic tree with reference and traded samples. Traded samples are indicated in the phylogram by their accession number.

**Fig. S9**. Maximum Likelihood rbcL phylogenetic tree with reference and traded samples. Traded samples are indicated in the phylogram by their accession number.

**Fig. S10**. Maximum Likelihood trnH-psbA phylogenetic tree with reference and traded samples. Traded samples are indicated in the phylogram by their accession number.

**Fig. S11**. Maximum Likelihood trnL phylogenetic tree with reference and traded samples. Traded samples are indicated in the phylogram by their accession number.

**Fig. S12.** Multispecies Coalescent (MSC) phylogenetic tree based on the nuclear loci dataset with the reference dataset only. The vertical lines on the right of the phylogeny represent supported clades with the associated species names.

**Fig. S13**. Analysis workflow of root samples in trade.

**Table S1**. Voucher specimens for the reference database. The herbaria where the reference material is deposited is indicated using Index Herbariorum abbreviations: BC for Institut Botànic de Barcelona, Spain; HUJ for Hebrew University, Israel; MA for Real Jardín Botánico, Spain; MARK for Cadi Ayyad University, Morocco; P for Muséum National d'Histoire Naturelle, France; O for Botanical Museum, University of Oslo, Norway; UPNA for Universidad Pública de Navarra; Spain; VIT for Museo de Ciencias Naturales de Alava, Spain.

**Table S2**. Voucher specimens for traded samples and their taxonomic identification. The first section of the table lists the samples identified from target capture using a phylogenetic framework, and the second section the samples identified using *blastn*.

**Table S3**. Reference sequences for the ITS tree with their respective article reference: [1](Guo, Ehrendorfer, & Samuel, 2004), [2](Ata, Abd El-Twab, Helmey, & Dahy, 2017), [3](Priya, Malik, & Babbar, 2018), [4](Oberprieler, 2004), [5](Sonboli, Stroka, Osaloo, & Oberprieler, 2012), [6](Ghorbani, Saeedi, & De Boer, 2017).

**Table S4**. Basic statistics on the number of reads per sample for the target sequencing and the genome skimming sequencing runs. The table is summarized in Figure S3.

**Table S5**. Average coverage, length of the matrices and missing data for the standard barcodes, ITS, the plastome and low copy nuclear markers.

**Table S6**. Percentage of adulteration and molecular identification of the different varieties of *A. pyrethrum* for each of the value chain stakeholders. This table is summarized in Figure 4.

**Table S7**. Sequencing recovery and identification success for the traded samples for each dataset. This table is summarized in Figure 1.

### SI Figures

**Figure S1.**

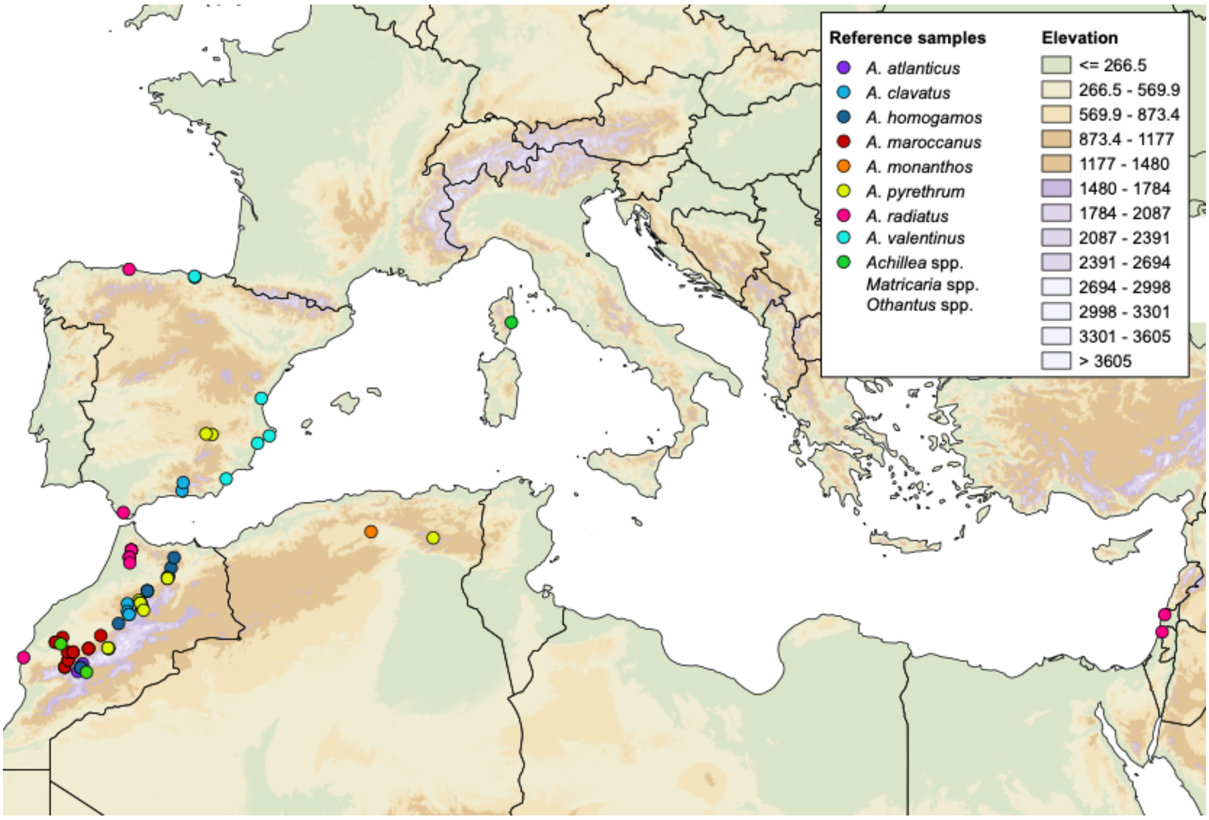

**Figure S2.**

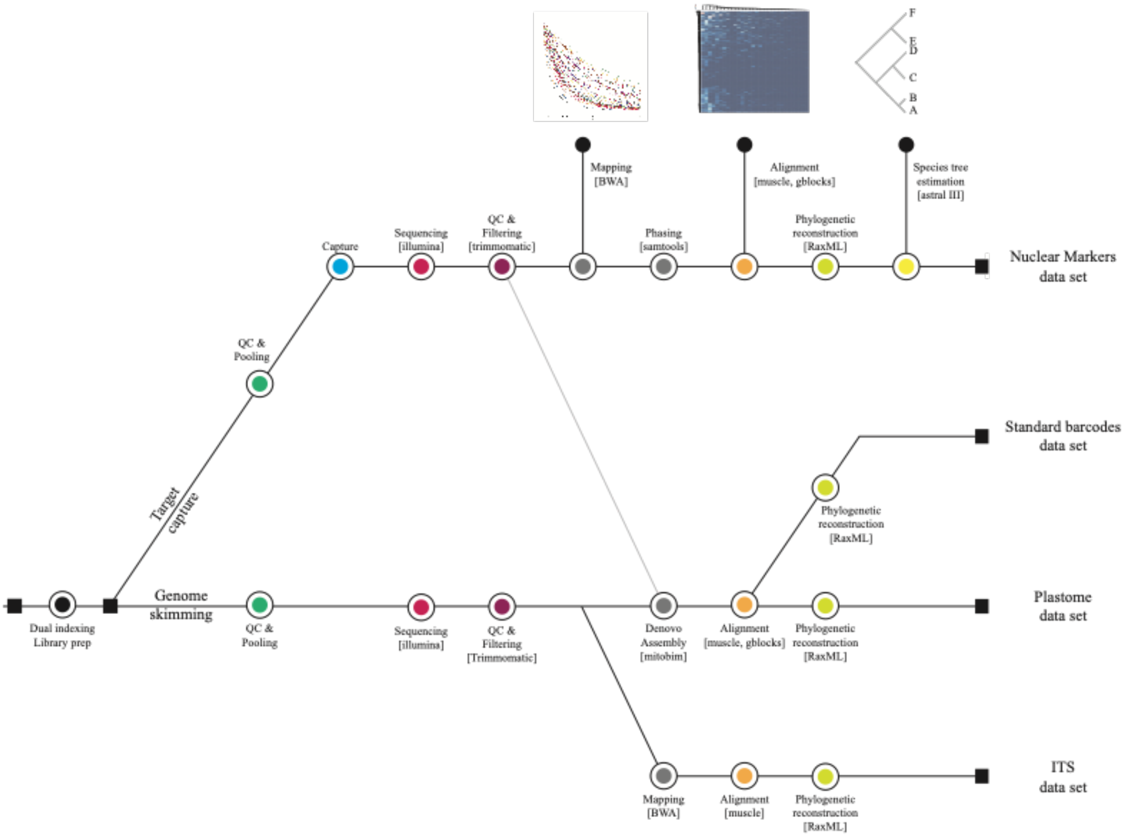
Mapping [BWA]
Alignment [muscle, gblocks]
Species tree estimation [astral III]
Capture
Sequencing [illumina]
QC & Filtering [trimmomatic]
Phasing [samtools]
Phylogenetic reconstruction [RaxML]
Nuclear Markers data set
QC & Pooling
Target capture
Standard barcodes data set
Phylogenetic reconstruction [RaxML]
Genome skimming
Dual indexing Library prep
QC & Pooling
Sequencing [illumina]
QC & Filtering [Trimmomatic]
Denovo Assembly [mitobim]
Alignment [muscle, gblocks]
Phylogenetic reconstruction [RaxML]
Plastome data set
Mapping [BWA]
Alignment [muscle]
Phylogenetic reconstruction [RaxML]
ITS data set

**Figure S3.**

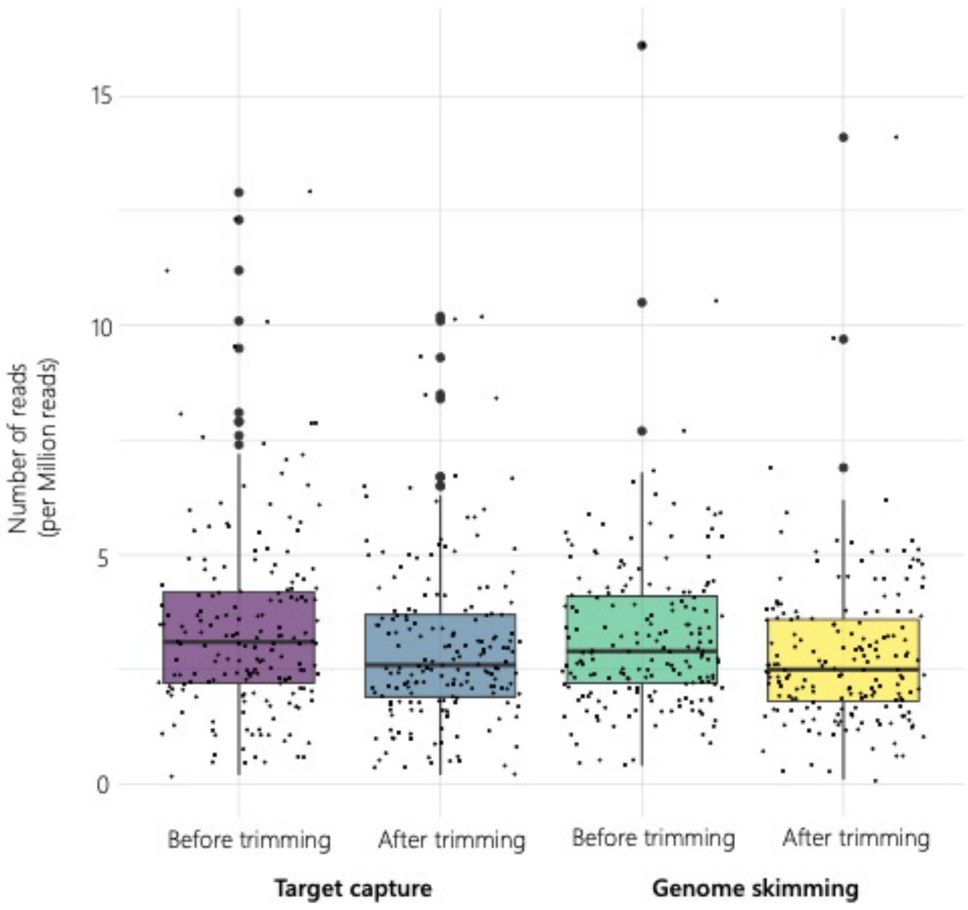
15
10
5
0
Number of reads
(per Million reads)
Before trimming
After trimming
Before trimming
After trimming
Target capture
Genome skimming

**Figure S4.**

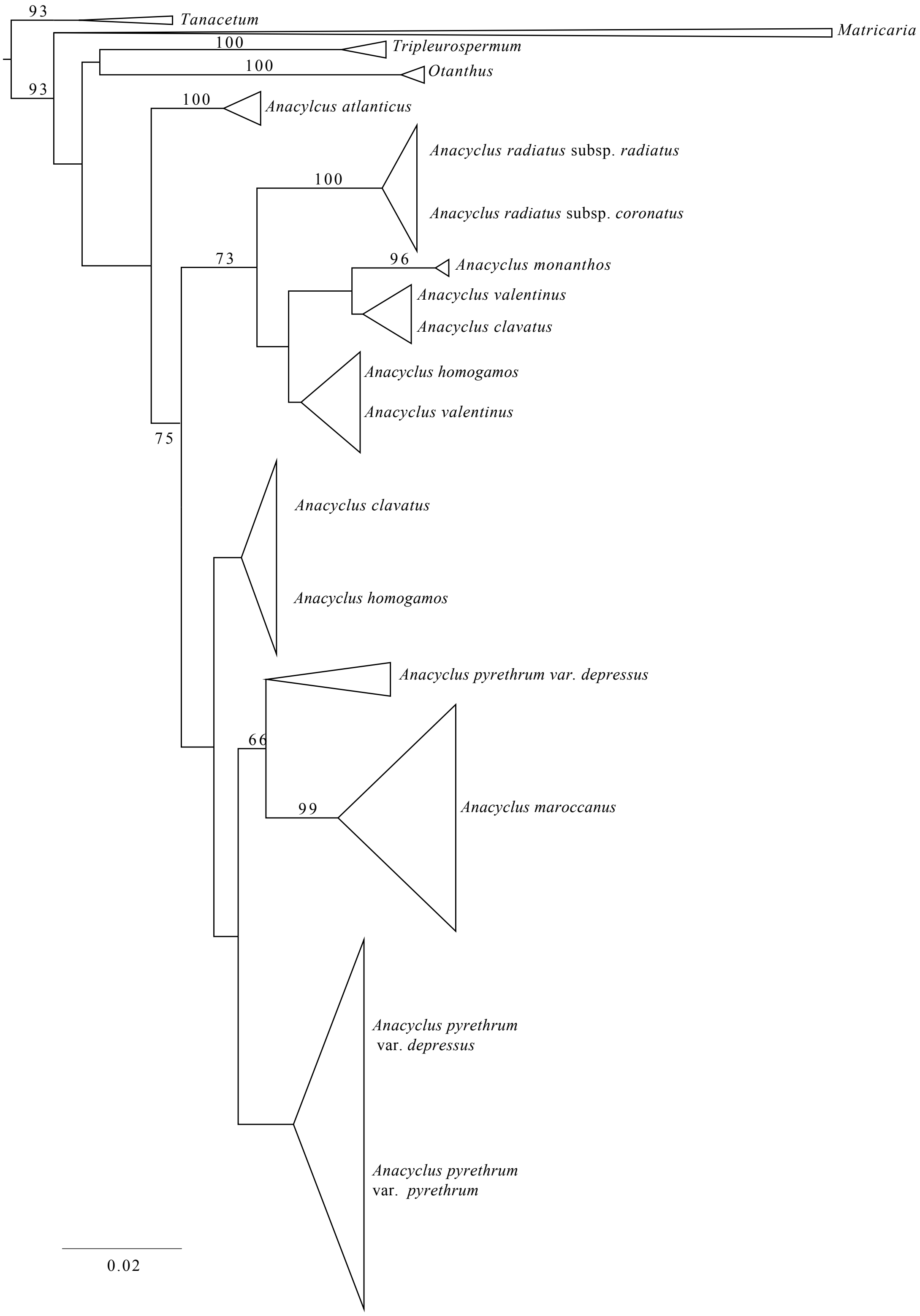
93
Tanacetum
Matricaria
100
Tripleurospermum
100
Otanthus
93
100
Anacylcus atlanticus
Anacyclus radiatus subsp. radiatus
100
Anacyclus radiatus subsp. coronatus
73
96
Anacyclus monanthos
Anacyclus valentinus
Anacyclus clavatus
Anacyclus homogamos
Anacyclus valentinus
75
Anacyclus clavatus
Anacyclus homogamos
Anacyclus pyrethrum var. depressus
66
99
Anacyclus maroccanus
Anacyclus pyrethrum var. depressus
Anacyclus pyrethrum var. pyrethrum
0.02

### Figure S5.

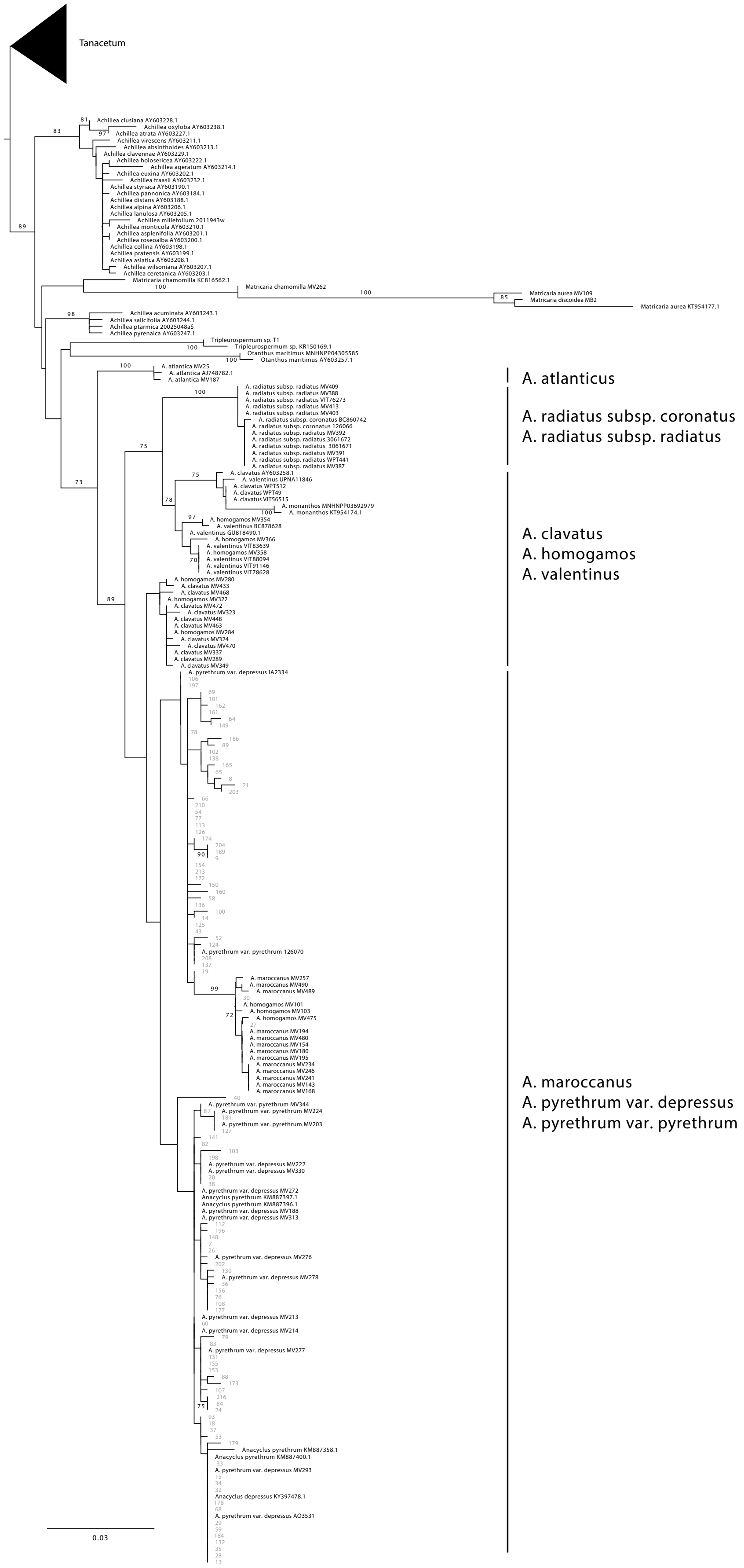

Tanacetum
Achillea clusiana AY603228.1
Achillea oxyloba AY603238.1
Achillea atrata AY603227.1
Achillea virescens AY603211.1
Achillea absinthoides AY603213.1
Achillea clavennae AY603229.1
Achillea holosericea AY603222.1
Achillea ageratum AY603214.1
Achillea euxina AY603202.1
Achillea fraasii AY603232.1
Achillea styriaca AY603190.1
Achillea pannonica AY603184.1
Achillea distans AY603188.1
Achillea alpina AY603206.1
Achillea lanulosa AY603205.1
Achillea millefolium 2011943w
Achillea monticola AY603210.1
Achillea asplenifolia AY603201.1
Achillea roseoalba AY603200.1
Achillea collina AY603198.1
Achillea pratensis AY603199.1
Achillea asiatica AY603208.1
Achillea wilsoniana AY603207.1
Achillea ceretanica AY603203.1
Matricaria chamomilla KC816562.1
Matricaria chamomilla MV262
Matricaria aurea MV109
Matricaria discoidea MB2
Matricaria aurea KT954177.1
Achillea acuminata AY603243.1
Achillea salicifolia AY603244.1
Achillea ptarmica 20025048aS
Achillea pyrenaica AY603247.1
Tripleurospermum sp. T1
Tripleurospermum sp. KR150169.1
Otanthus maritimus MNHNPP04305585
Otanthus maritimus AY603257.1
A. atlantica MV25
A. atlantica AJ748782.1
A. atlantica MV187
A. atlanticus
A. radiatus subsp. coronatus
A. radiatus subsp. radiatus
A. clavatus
A. homogamos
A. valentinus
A. maroccanus
A. pyrethrum var. depressus
A. pyrethrum var. pyrethrum
0.03

**Figure S6.**

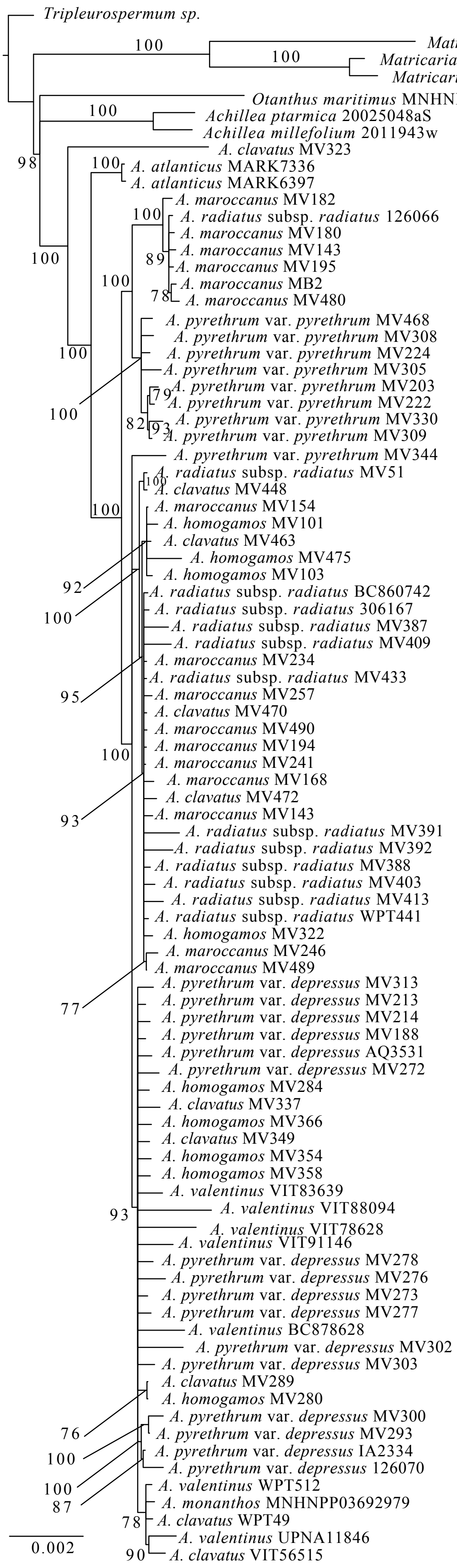
Tripleurospermum sp.
Matricaria discoidea MB2
Matricaria chamomilla MV262
Matricaria aurea MV109
Otanthus maritimus MNHNPP04305585
Achillea ptarmica 20025048aS
Achillea millefolium 2011943w
A. clavatus MV323
A. atlanticus MARK7336
A. atlanticus MARK6397
A. maroccanus MV182
A. radiatus subsp. radiatus 126066
A. maroccanus MV180
A. maroccanus MV143
A. maroccanus MV195
A. maroccanus MB2
A. maroccanus MV480
A. pyrethrum var. pyrethrum MV468
A. pyrethrum var. pyrethrum MV308
A. pyrethrum var. pyrethrum MV224
A. pyrethrum var. pyrethrum MV305
A. pyrethrum var. pyrethrum MV203
A. pyrethrum var. pyrethrum MV222
A. pyrethrum var. pyrethrum MV330
A. pyrethrum var. pyrethrum MV309
A. pyrethrum var. pyrethrum MV344
A. radiatus subsp. radiatus MV51
A. clavatus MV448
A. maroccanus MV154
A. homogamos MV101
A. clavatus MV463
A. homogamos MV475
A. homogamos MV103
A. radiatus subsp. radiatus BC860742
A. radiatus subsp. radiatus 306167
A. radiatus subsp. radiatus MV387
A. radiatus subsp. radiatus MV409
A. maroccanus MV234
A. radiatus subsp. radiatus MV433
A. maroccanus MV257
A. clavatus MV470
A. maroccanus MV490
A. maroccanus MV194
A. maroccanus MV241
A. maroccanus MV168
A. clavatus MV472
A. maroccanus MV143
A. radiatus subsp. radiatus MV391
A. radiatus subsp. radiatus MV392
A. radiatus subsp. radiatus MV388
A. radiatus subsp. radiatus MV403
A. radiatus subsp. radiatus MV413
A. radiatus subsp. radiatus WPT441
A. homogamos MV322
A. maroccanus MV246
A. maroccanus MV489
A. pyrethrum var. depressus MV313
A. pyrethrum var. depressus MV213
A. pyrethrum var. depressus MV214
A. pyrethrum var. depressus MV188
A. pyrethrum var. depressus AQ3531
A. pyrethrum var. depressus MV272
A. homogamos MV284
A. clavatus MV337
A. homogamos MV366
A. clavatus MV349
A. homogamos MV354
A. homogamos MV358
A. valentinus VIT83639
A. valentinus VIT88094
A. valentinus VIT78628
A. valentinus VIT91146
A. pyrethrum var. depressus MV278
A. pyrethrum var. depressus MV276
A. pyrethrum var. depressus MV273
A. pyrethrum var. depressus MV277
A. valentinus BC878628
A. pyrethrum var. depressus MV302
A. pyrethrum var. depressus MV303
A. clavatus MV289
A. homogamos MV280
A. pyrethrum var. depressus MV300
A. pyrethrum var. depressus MV293
A. pyrethrum var. depressus IA2334
A. pyrethrum var. depressus 126070
A. valentinus WPT512
A. monanthos MNHNPP03692979
A. clavatus WPT49
A. valentinus UPNA11846
A. clavatus VIT56515
0.002

#### Figure S7.

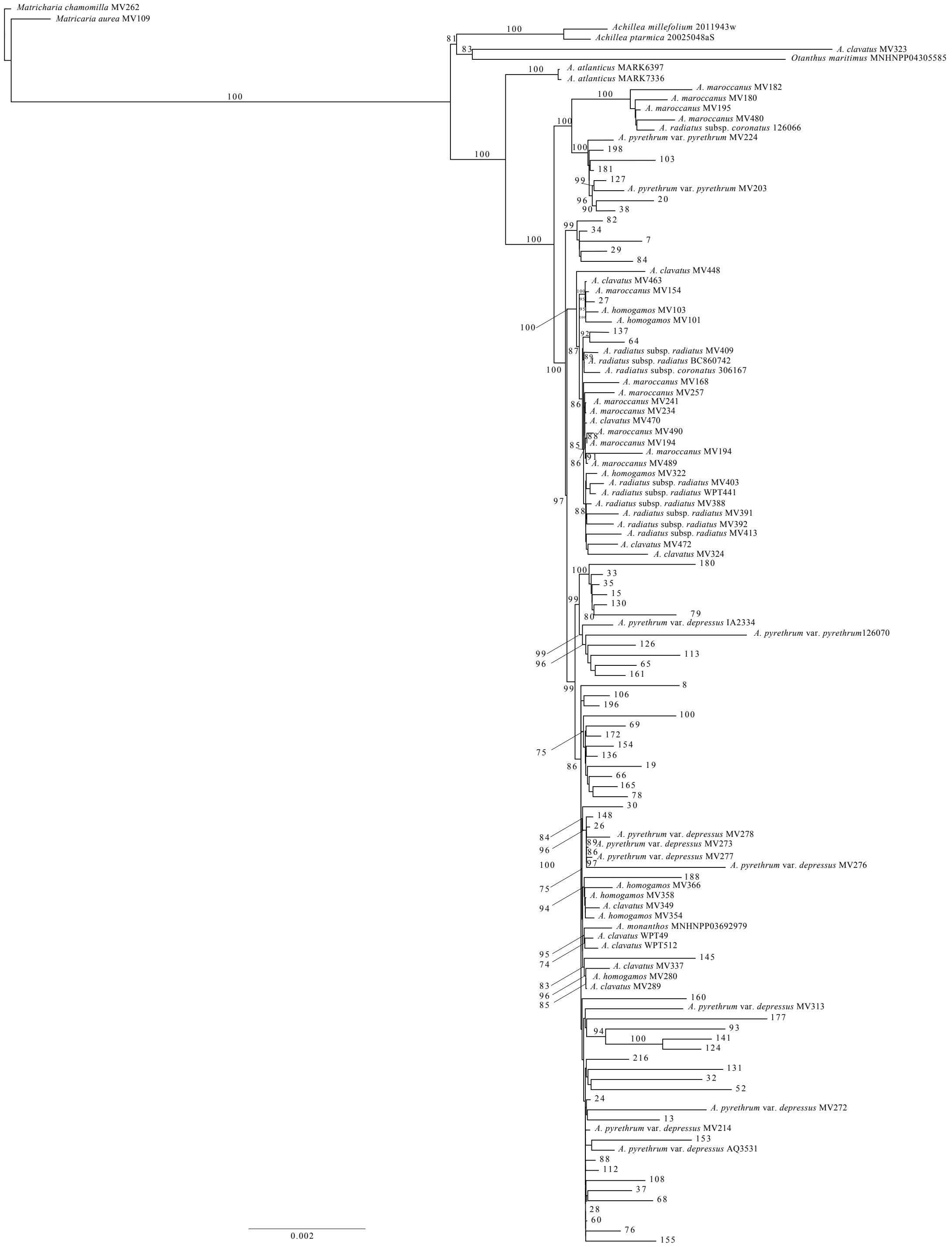
Matricharia chamomilla MV262
Matricaria aurea MV109
Achillea millefolium 2011943w
Achillea ptarmica 20025048aS
A. clavatus MV323
Otanthus maritimus MNHNPP04305585
A. atlanticus MARK6397
A. atlanticus MARK7336
A. maroccanus MV182
A. maroccanus MV180
A. maroccanus MV195
A. maroccanus MV480
A. radiatus subsp. coronatus 126066
A. pyrethrum var. pyrethrum MV224
198
103
181
127
A. pyrethrum var. pyrethrum MV203
20
38
82
34
7
29
84
A. clavatus MV448
A. clavatus MV463
A. maroccanus MV154
27
A. homogamos MV103
A. homogamos MV101
137
64
A. radiatus subsp. radiatus MV409
A. radiatus subsp. radiatus BC860742
A. radiatus subsp. coronatus 306167
A. maroccanus MV168
A. maroccanus MV257
A. maroccanus MV241
A. maroccanus MV234
A. clavatus MV470
A. maroccanus MV490
A. maroccanus MV194
A. maroccanus MV194
A. maroccanus MV489
A. homogamos MV322
A. radiatus subsp. radiatus MV403
A. radiatus subsp. radiatus WPT441
A. radiatus subsp. radiatus MV388
A. radiatus subsp. radiatus MV391
A. radiatus subsp. radiatus MV392
A. radiatus subsp. radiatus MV413
A. clavatus MV472
A. clavatus MV324
180
33
35
15
130
79
A. pyrethrum var. depressus IA2334
A. pyrethrum var. pyrethrum126070
126
113
65
161
8
106
196
100
69
172
154
136
19
66
165
78
30
148
26
A. pyrethrum var. depressus MV278
A. pyrethrum var. depressus MV273
A. pyrethrum var. depressus MV277
A. pyrethrum var. depressus MV276
188
A. homogamos MV366
A. homogamos MV358
A. clavatus MV349
A. homogamos MV354
A. monanthos MNHNPP03692979
A. clavatus WPT49
A. clavatus WPT512
145
A. clavatus MV337
A. homogamos MV280
A. clavatus MV289
160
A. pyrethrum var. depressus MV313
177
93
141
124
216
131
32
52
24
A. pyrethrum var. depressus MV272
13
A. pyrethrum var. depressus MV214
153
A. pyrethrum var. depressus AQ3531
88
112
108
37
68
28
60
76
155
0.002

**Figure S8.**

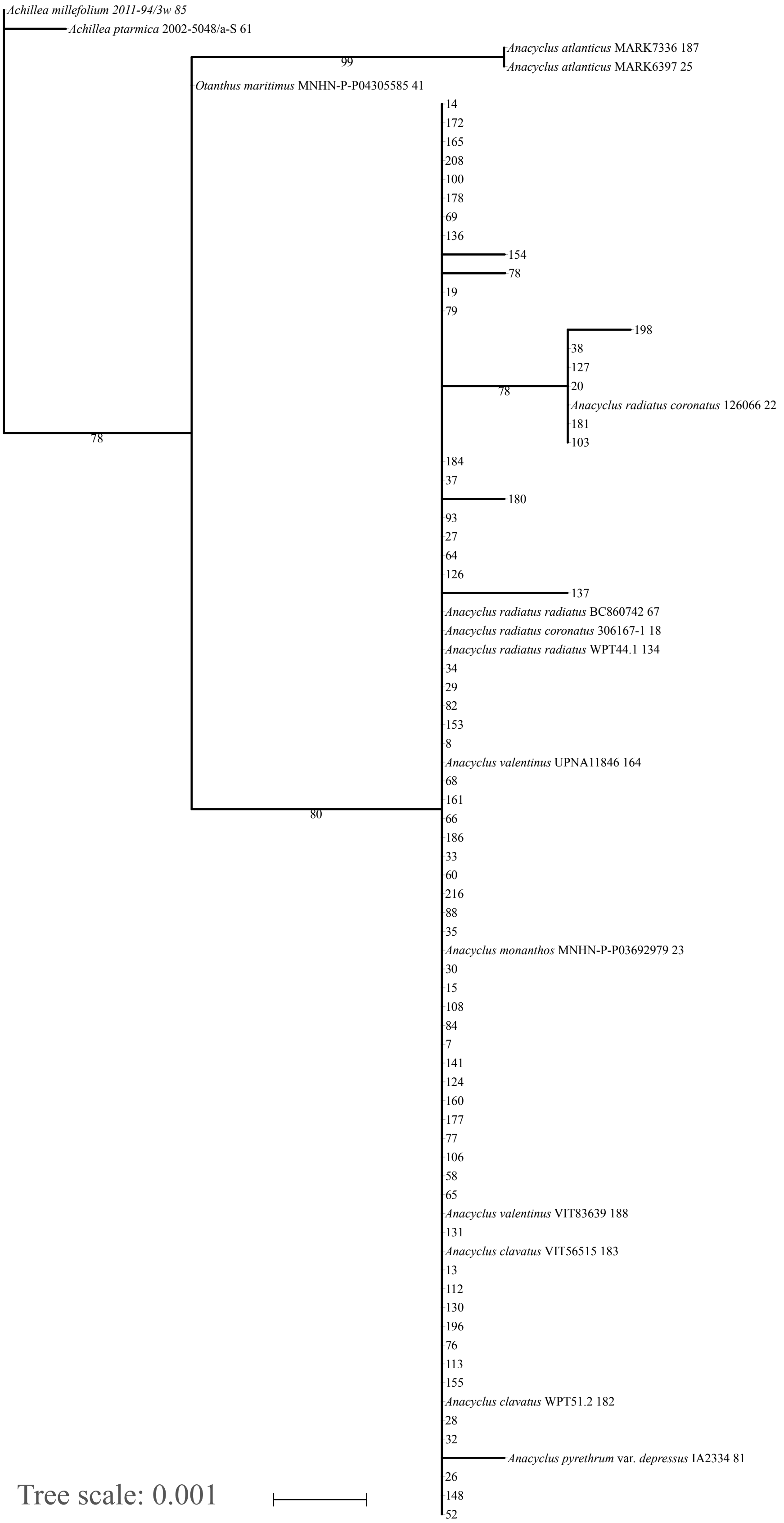
Achillea millefolium 2011-94/3w 85
Achillea ptarmica 2002-5048/a-S 61
Anacyclus atlanticus MARK7336 187
Anacyclus atlanticus MARK6397 25
99
Otanthus maritimus MNHN-P-P04305585 41
14
172
165
208
100
178
69
136
154
78
19
79
198
38
127
20
78
Anacyclus radiatus coronatus 126066 22
181
103
78
184
37
180
93
27
64
126
137
Anacyclus radiatus radiatus BC860742 67
Anacyclus radiatus coronatus 306167-1 18
Anacyclus radiatus radiatus WPT44.1 134
34
29
82
153
8
Anacyclus valentinus UPNA11846 164
68
161
80
66
186
33
60
216
88
35
Anacyclus monanthos MNHN-P-P03692979 23
30
15
108
84
7
141
124
160
177
77
106
58
65
Anacyclus valentinus VIT83639 188
131
Anacyclus clavatus VIT56515 183
13
112
130
196
76
113
155
Anacyclus clavatus WPT51.2 182
28
32
Anacyclus pyrethrum var. depressus IA2334 81
26
148
52
Tree scale: 0.001

**Figure S9.**

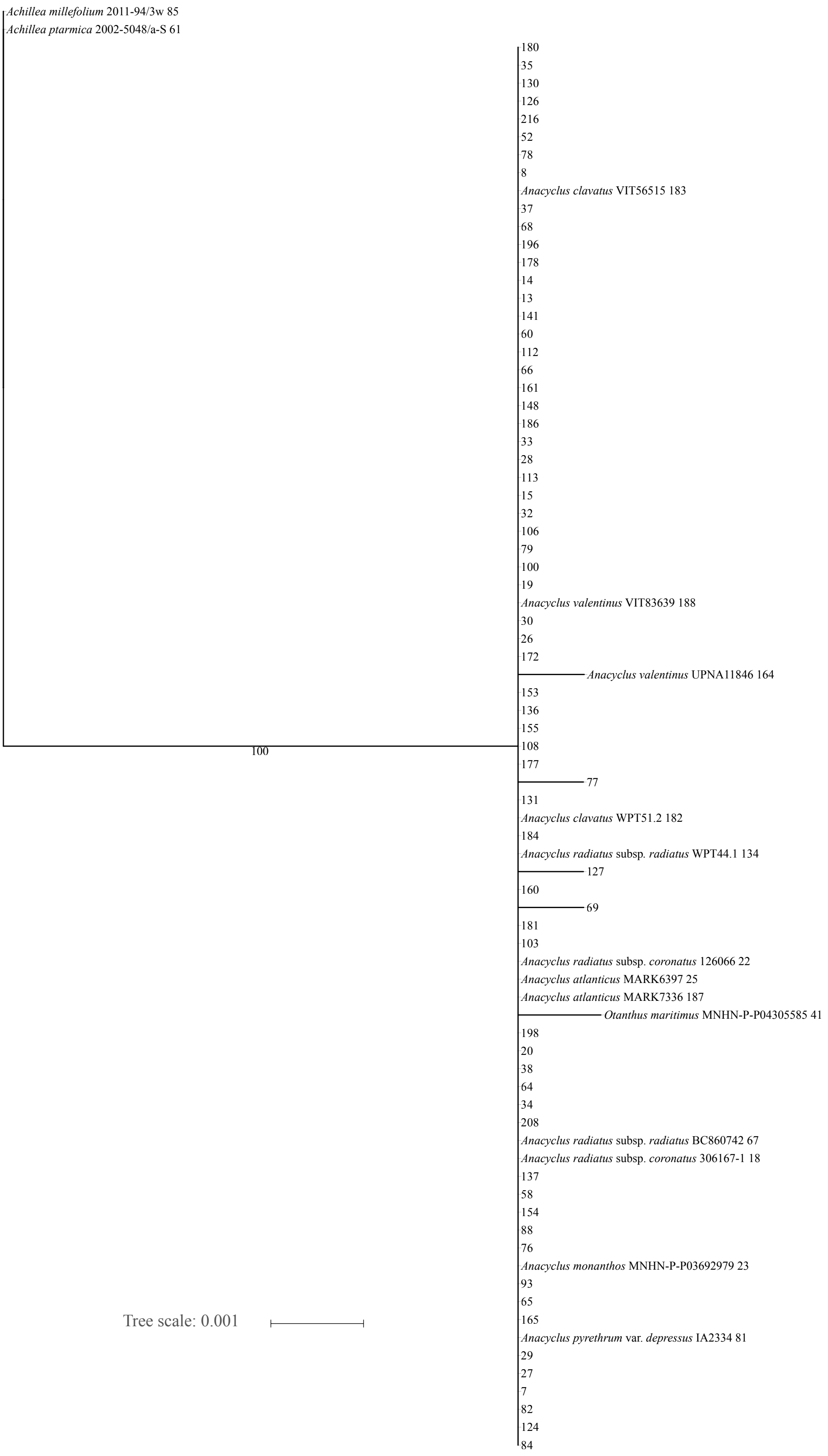
Achillea millefolium 2011-94/3w 85
Achillea ptarmica 2002-5048/a-S 61
180
35
130
126
216
52
78
8
Anacyclus clavatus VIT56515 183
37
68
196
178
14
13
141
60
112
66
161
148
186
33
28
113
15
32
106
79
100
19
Anacyclus valentinus VIT83639 188
30
26
172
Anacyclus valentinus UPNA11846 164
153
136
155
108
177
77
131
Anacyclus clavatus WPT51.2 182
184
Anacyclus radiatus subsp. radiatus WPT44.1 134
127
160
69
181
103
Anacyclus radiatus subsp. coronatus 126066 22
Anacyclus atlanticus MARK6397 25
Anacyclus atlanticus MARK7336 187
Otanthus maritimus MNHN-P-P04305585 41
198
20
38
64
34
208
Anacyclus radiatus subsp. radiatus BC860742 67
Anacyclus radiatus subsp. coronatus 306167-1 18
137
58
154
88
76
Anacyclus monanthos MNHN-P-P03692979 23
93
65
165
Anacyclus pyrethrum var. depressus IA2334 81
29
27
7
82
124
84
100
Tree scale: 0.001

**Figure S10.**

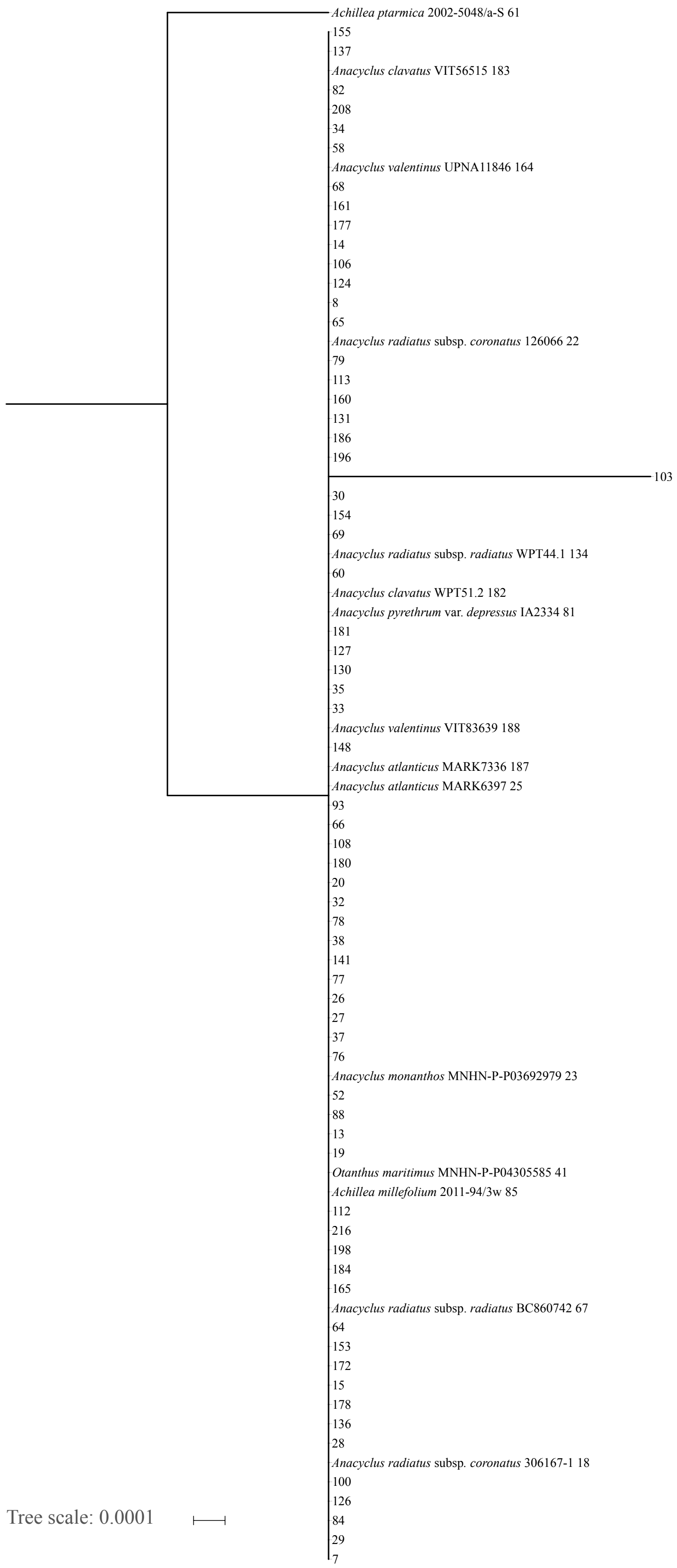
Achillea ptarmica 2002-5048/a-S 61
155
137
Anacyclus clavatus VIT56515 183
82
208
34
58
Anacyclus valentinus UPNA11846 164
68
161
177
14
106
124
8
65
Anacyclus radiatus subsp. coronatus 126066 22
79
113
160
131
186
196
103
30
154
69
Anacyclus radiatus subsp. radiatus WPT44.1 134
60
Anacyclus clavatus WPT51.2 182
Anacyclus pyrethrum var. depressus IA2334 81
181
127
130
35
33
Anacyclus valentinus VIT83639 188
148
Anacyclus atlanticus MARK7336 187
Anacyclus atlanticus MARK6397 25
93
66
108
180
20
32
78
38
141
77
26
27
37
76
Anacyclus monanthos MNHN-P-P03692979 23
52
88
13
19
Otanthus maritimus MNHN-P-P04305585 41
Achillea millefolium 2011-94/3w 85
112
216
198
184
165
Anacyclus radiatus subsp. radiatus BC860742 67
64
153
172
15
178
136
28
Anacyclus radiatus subsp. coronatus 306167-1 18
100
126
84
29
7
Tree scale: 0.0001

#### Figure S11.

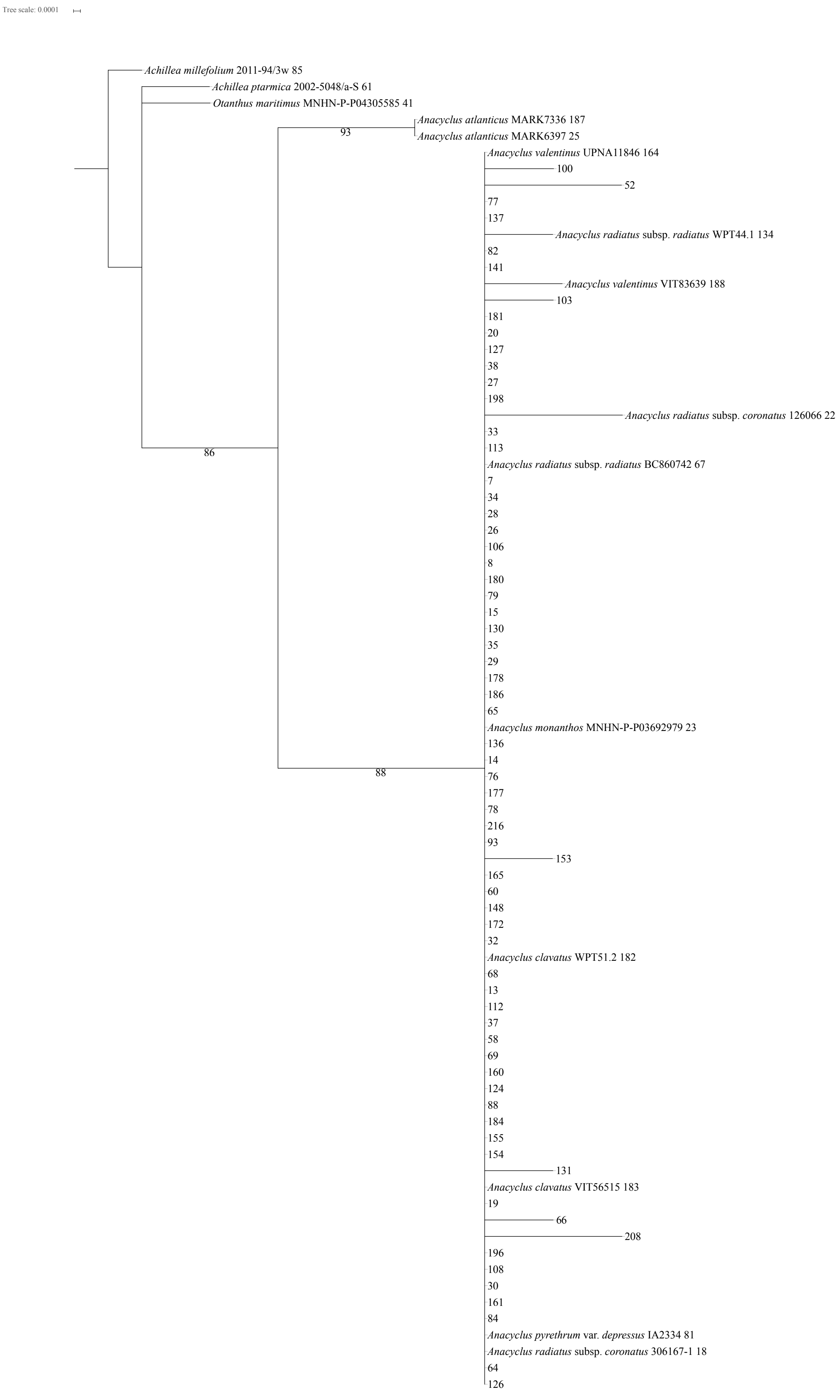
Tree scale: 0.0001
Achillea millefolium 2011-94/3w 85
Achillea ptarmica 2002-5048/a-S 61
Otanthus maritimus MNHN-P-P04305585 41
Anacyclus atlanticus MARK7336 187
93
Anacyclus atlanticus MARK6397 25
Anacyclus valentinus UPNA11846 164
100
52
77
137
Anacyclus radiatus subsp. radiatus WPT44.1 134
82
141
Anacyclus valentinus VIT83639 188
103
181
20
127
38
27
198
Anacyclus radiatus subsp. coronatus 126066 22
33
113
86
Anacyclus radiatus subsp. radiatus BC860742 67
7
34
28
26
106
8
180
79
15
130
35
29
178
186
65
Anacyclus monanthos MNHN-P-P03692979 23
136
14
88
76
177
78
216
93
153
165
60
148
172
32
Anacyclus clavatus WPT51.2 182
68
13
112
37
58
69
160
124
88
184
155
154
131
Anacyclus clavatus VIT56515 183
19
66
208
196
108
30
161
84
Anacyclus pyrethrum var. depressus IA2334 81
Anacyclus radiatus subsp. coronatus 306167-1 18
64
126

**Figure S12.**

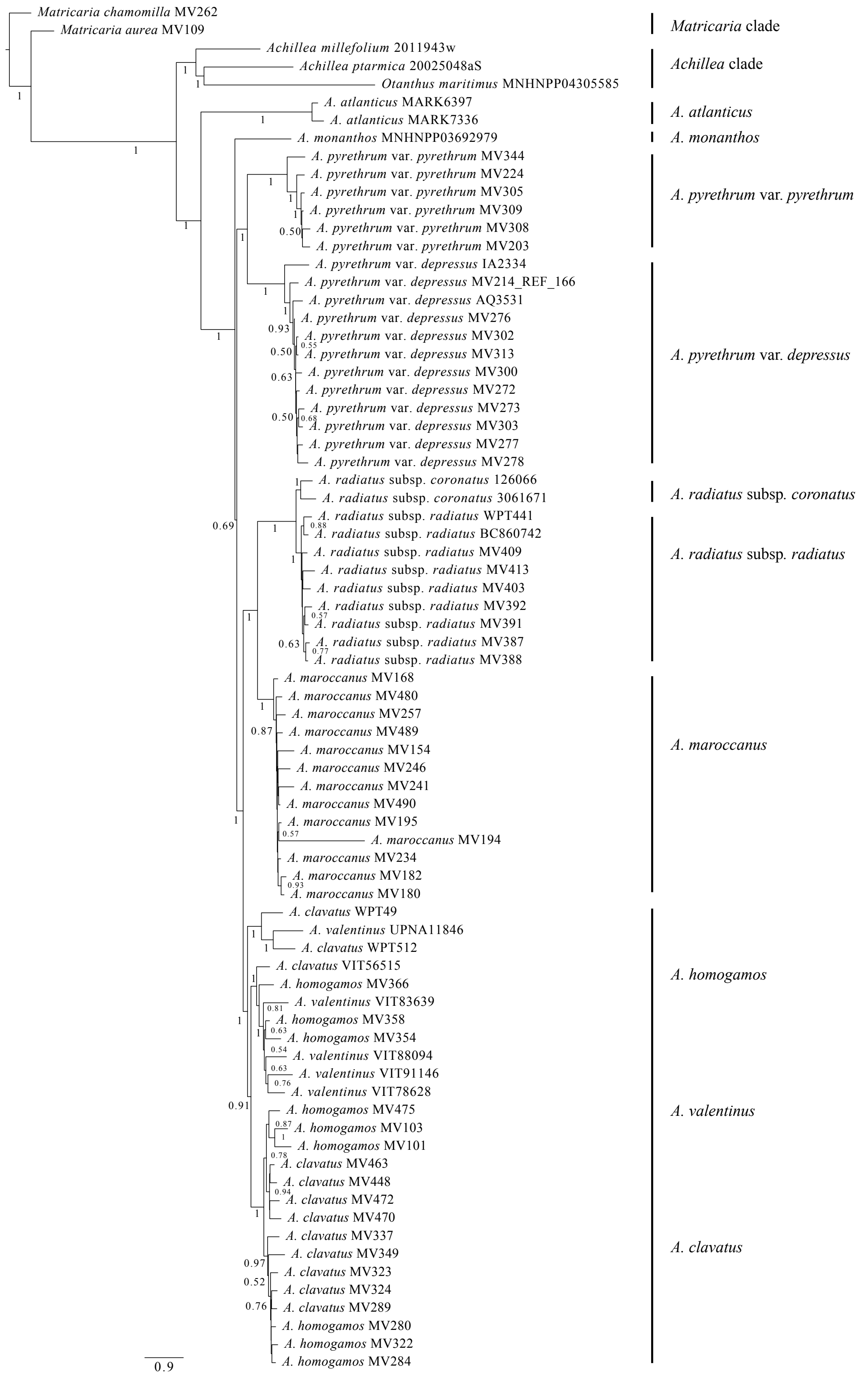
Matricaria chamomilla MV262
Matricaria aurea MV109
Achillea millefolium 2011943w
Achillea ptarmica 20025048aS
Otanthus maritimus MNHNPP04305585
A. atlanticus MARK6397
A. atlanticus MARK7336
A. monanthos MNHNPP03692979
A. pyrethrum var. pyrethrum MV344
A. pyrethrum var. pyrethrum MV224
A. pyrethrum var. pyrethrum MV305
A. pyrethrum var. pyrethrum MV309
A. pyrethrum var. pyrethrum MV308
A. pyrethrum var. pyrethrum MV203
A. pyrethrum var. depressus IA2334
A. pyrethrum var. depressus MV214_REF_166
A. pyrethrum var. depressus AQ3531
A. pyrethrum var. depressus MV276
A. pyrethrum var. depressus MV302
A. pyrethrum var. depressus MV313
A. pyrethrum var. depressus MV300
A. pyrethrum var. depressus MV272
A. pyrethrum var. depressus MV273
A. pyrethrum var. depressus MV303
A. pyrethrum var. depressus MV277
A. pyrethrum var. depressus MV278
A. radiatus subsp. coronatus 126066
A. radiatus subsp. coronatus 3061671
A. radiatus subsp. radiatus WPT441
A. radiatus subsp. radiatus BC860742
A. radiatus subsp. radiatus MV409
A. radiatus subsp. radiatus MV413
A. radiatus subsp. radiatus MV403
A. radiatus subsp. radiatus MV392
A. radiatus subsp. radiatus MV391
A. radiatus subsp. radiatus MV387
A. radiatus subsp. radiatus MV388
A. maroccanus MV168
A. maroccanus MV480
A. maroccanus MV257
A. maroccanus MV489
A. maroccanus MV154
A. maroccanus MV246
A. maroccanus MV241
A. maroccanus MV490
A. maroccanus MV195
A. maroccanus MV194
A. maroccanus MV234
A. maroccanus MV182
A. maroccanus MV180
A. clavatus WPT49
A. valentinus UPNA11846
A. clavatus WPT512
A. clavatus VIT56515
A. homogamos MV366
A. valentinus VIT83639
A. homogamos MV358
A. homogamos MV354
A. valentinus VIT88094
A. valentinus VIT91146
A. valentinus VIT78628
A. homogamos MV475
A. homogamos MV103
A. homogamos MV101
A. clavatus MV463
A. clavatus MV448
A. clavatus MV472
A. clavatus MV470
A. clavatus MV337
A. clavatus MV349
A. clavatus MV323
A. clavatus MV324
A. clavatus MV289
A. homogamos MV280
A. homogamos MV322
A. homogamos MV284
0.9
Matricaria clade
Achillea clade
A. atlanticus
A. monanthos
A. pyrethrum var. pyrethrum
A. pyrethrum var. depressus
A. radiatus subsp. coronatus
A. radiatus subsp. radiatus
A. maroccanus
A. homogamos
A. valentinus
A. clavatus

**Figure S13.**

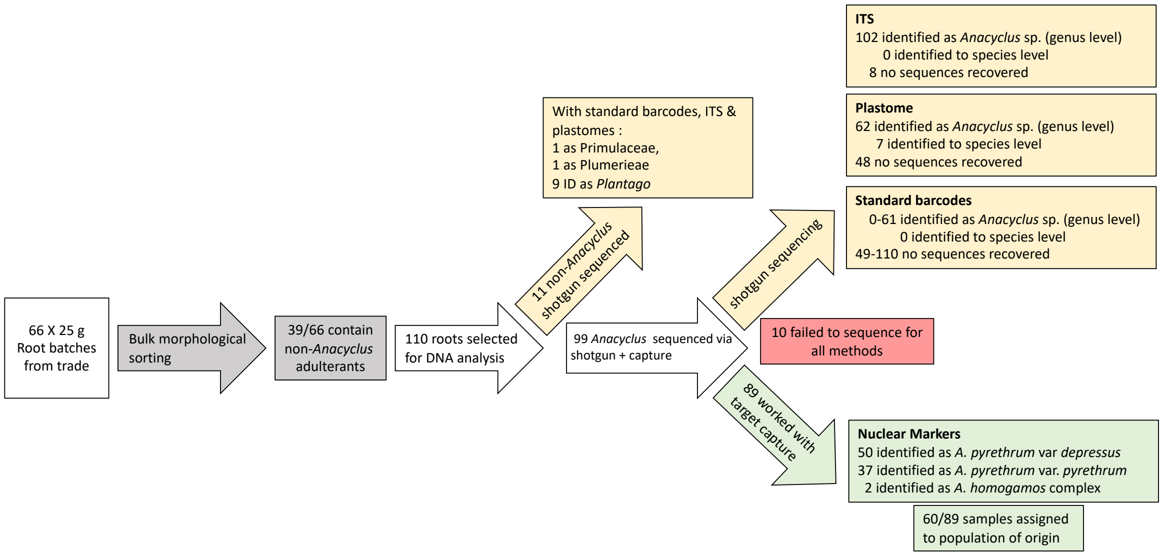
66 X 25 g
Root batches
from trade
Bulk morphological sorting
39/66 contain non-*Anacyclus* adulterants
110 roots selected for DNA analysis
11 non-*Anacyclus* shotgun sequenced
With standard barcodes, ITS & plastomes :
1 as Primulaceae,
1 as Plumerieae
9 ID as *Plantago*
99 *Anacyclus* sequenced via shotgun + capture
shotgun sequencing
ITS
102 identified as *Anacyclus* sp. (genus level)
0 identified to species level
8 no sequences recovered
Plastome
62 identified as *Anacyclus* sp. (genus level)
7 identified to species level
48 no sequences recovered
Standard barcodes
0-61 identified as *Anacyclus* sp. (genus level)
0 identified to species level
49-110 no sequences recovered
10 failed to sequence for all methods
89 worked with target capture
Nuclear Markers
50 identified as *A. pyrethrum* var *depressus*
37 identified as *A. pyrethrum* var. *pyrethrum*
2 identified as *A. homogamos* complex
60/89 samples assigned to population of origin

#### SI Tables

#### Table S1.

| Genus | Species | subsp. / var. | Collection ID | Index Herbariorum | ID during sequencing | Date collected | GPS | | Country |
|---|---|---|---|---|---|---|---|---|---|
| | | | | | | | ° N | ° E | |
| *Achillea* | *millefolium* | | 2011-94/3w | O | 85 | 2015 | 59,92 | 10,77 | Norway |
| *Achillea* | *ptarmica* | | 2002-5048/a-S | O | 61 | 2015 | 59,92 | 10,77 | Norway |
| *Anacyclus* | *atlanticus* | | MARK6397 | MARK | 25 | 16/07/2008 | 31,03 | -7,57 | Morocco |
| *Anacyclus* | *atlanticus* | | MARK7336 | MARK | 187 | 2005 | 31,28 | -7,38 | Morocco |
| *Anacyclus* | *clavatus* | | MV289 | O | 121 | 18/04/2015 | 33,16 | -5,07 | Morocco |
| *Anacyclus* | *clavatus* | | MV323 | O | 171 | 18/04/2015 | 33,70 | -4,84 | Morocco |
| *Anacyclus* | *clavatus* | | MV324 | O | 143 | 18/04/2015 | 33,70 | -4,84 | Morocco |
| *Anacyclus* | *clavatus* | | MV337 | O | 145 | 19/04/2015 | 34,17 | -4,01 | Morocco |
| *Anacyclus* | *clavatus* | | MV349 | O | 104 | 19/04/2015 | 34,11 | -4,05 | Morocco |
| *Anacyclus* | *clavatus* | | MV448 | O | 95 | 22/04/2015 | 33,28 | -5,62 | Morocco |
| *Anacyclus* | *clavatus* | | MV463 | O | 122 | 22/04/2015 | 33,03 | -5,62 | Morocco |
| *Anacyclus* | *clavatus* | | MV470 | O | 168 | 23/04/2015 | 32,93 | -5,56 | Morocco |
| *Anacyclus* | *clavatus* | | MV472 | O | 86 | 23/04/2015 | 32,93 | -5,56 | Morocco |
| *Anacyclus* | *clavatus* | | VIT 56515 | VIT | 183 | 1997 | 43,30 | -2,98 | Spain |
| *Anacyclus* | *clavatus* | | WPT49.1 | O | 158 | 23/06/2015 | 36,91 | -3,48 | Spain |
| *Anacyclus* | *clavatus* | | WPT51.2 | O | 182 | 23/06/2015 | 37,16 | -3,44 | Spain |
| *Anacyclus* | *homogamos* | | MV101 | O | 176 | 04/04/2015 | 31,16 | -7,46 | Morocco |
| *Anacyclus* | *homogamos* | | MV103 | O | 200 | 04/04/2015 | 31,16 | -7,46 | Morocco |
| *Anacyclus* | *homogamos* | | MV280 | O | 75 | 18/04/2015 | 33,22 | -5,06 | Morocco |
| *Anacyclus* | *homogamos* | | MV284 | O | 170 | 18/04/2015 | 33,16 | -5,07 | Morocco |
| *Anacyclus* | *homogamos* | | MV322 | O | 48 | 18/04/2015 | 33,70 | -4,84 | Morocco |
| *Anacyclus* | *homogamos* | | MV354 | O | 71 | 19/04/2015 | 34,45 | -3,91 | Morocco |
| *Anacyclus* | *homogamos* | | MV358 | O | 73 | 19/04/2015 | 34,78 | -3,79 | Morocco |
| *Anacyclus* | *homogamos* | | MV366 | O | 2 | 20/04/2015 | 34,14 | -4,06 | Morocco |
| *Anacyclus* | *homogamos* | | MV475 | O | 194 | 23/04/2015 | 32,63 | -5,97 | Morocco |
| *Anacyclus* | *maroccanus* | | MV154 | O | 175 | 09/04/2015 | 31,17 | -8,08 | Morocco |
| *Anacyclus* | *maroccanus* | | MV168 | O | 6 | 09/04/2015 | 31,18 | -8,09 | Morocco |
| *Anacyclus* | *maroccanus* | | MV180 | O | 123 | 10/04/2015 | 31,41 | -7,95 | Morocco |
| *Anacyclus* | *maroccanus* | | MV182 | O | 1 | 10/04/2015 | 31,41 | -7,95 | Morocco |
| *Anacyclus* | *maroccanus* | | MV194 | O | 94 | 11/04/2015 | 31,80 | -7,15 | Morocco |
| *Anacyclus* | *maroccanus* | | MV195 | O | 118 | 11/04/2015 | 31,80 | -7,15 | Morocco |
| *Anacyclus* | *maroccanus* | | MV234 | O | 199 | 12/04/2015 | 32,01 | -8,45 | Morocco |
| *Anacyclus* | *maroccanus* | | MV241 | O | 11 | 12/04/2015 | 31,69 | -7,98 | Morocco |
| *Anacyclus* | *maroccanus* | | MV246 | O | 56 | 12/04/2015 | 32,17 | -8,17 | Morocco |
| *Anacyclus* | *maroccanus* | | MV257 | O | 144 | 12/04/2015 | 32,01 | -8,45 | Morocco |
| *Anacyclus* | *maroccanus* | | MV480 | O | 169 | 23/04/2015 | 32,23 | -6,67 | Morocco |
| *Anacyclus* | *maroccanus* | | MV489 | O | 193 | 23/04/2015 | 31,68 | -7,76 | Morocco |
| *Anacyclus* | *maroccanus* | | MV490 | O | 50 | 23/04/2015 | 31,68 | -7,76 | Morocco |
| *Anacyclus* | *monanthos* | | MNHN-P-P03692979 | P | 23 | 28/04/1990 | 35,61 | 3,94 | Algeria |
| *Anacyclus* | *pyrethrum* | *depressus* | AQ3531 | MA | 109 | 07/04/2015 | 38,65 | -2,31 | Spain |
| *Anacyclus* | *pyrethrum* | *depressus* | IA2334 | MA | 81 | 07/04/2015 | 34,18 | -2,54 | Morocco |
| *Anacyclus* | *pyrethrum* | *depressus* | MV214 | O | 166 | 11/04/2015 | 31,79 | -4,09 | Morocco |
| *Anacyclus* | *pyrethrum* | *depressus* | MV272 | O | 12 | 18/04/2015 | 33,39 | -5,17 | Morocco |
| *Anacyclus* | *pyrethrum* | *depressus* | MV273 | O | 201 | 18/04/2015 | 33,31 | -5,11 | Morocco |
| *Anacyclus* | *pyrethrum* | *depressus* | MV276 | O | 10 | 18/04/2015 | 33,31 | -5,11 | Morocco |
| *Anacyclus* | *pyrethrum* | *depressus* | MV277 | O | 55 | 18/04/2015 | 33,31 | -5,11 | Morocco |
| *Anacyclus* | *pyrethrum* | *depressus* | MV278 | O | 129 | 18/04/2015 | 33,31 | -5,11 | Morocco |
| *Anacyclus* | *pyrethrum* | *depressus* | MV313 | O | 167 | 18/04/2015 | 33,07 | -4,99 | Morocco |
| *Anacyclus* | *pyrethrum* | *pyrethrum* | 126070 | HUJ | 206 | 20/04/2015 | 35,41 | 6,38 | Algeria |
| *Anacyclus* | *pyrethrum* | *pyrethrum* | MV203 | O | 47 | 11/04/2015 | 31,82 | -6,36 | Morocco |
| *Anacyclus* | *pyrethrum* | *pyrethrum* | MV224 | O | 3 | 11/04/2015 | 31,82 | -6,39 | Morocco |
| *Anacyclus* | *pyrethrum* | *pyrethrum* | MV344 | O | 146 | 19/04/2015 | 34,11 | -4,05 | Morocco |
| *Anacyclus* | *radiatus* | *coronatus* | 126066 | HUJ | 22 | 2001 | 32,92 | 35,09 | Israel |
| *Anacyclus* | *radiatus* | *coronatus* | 306167-1 | HUJ | 18 | 2000 | 32,35 | 34,99 | Israel |
| *Anacyclus* | *radiatus* | *radiatus* | BC860742 | BC | 67 | 2009 | 43,53 | -5,56 | Spain |
| *Anacyclus* | *radiatus* | *radiatus* | MV387 | O | 4 | 22/04/2015 | 35,02 | -5,47 | Morocco |
| *Anacyclus* | *radiatus* | *radiatus* | MV388 | O | 5 | 22/04/2015 | 35,02 | -5,47 | Morocco |
| *Anacyclus* | *radiatus* | *radiatus* | MV391 | O | 128 | 22/04/2015 | 35,02 | -5,47 | Morocco |
| *Anacyclus* | *radiatus* | *radiatus* | MV392 | O | 205 | 22/04/2015 | 35,02 | -5,47 | Morocco |
| *Anacyclus* | *radiatus* | *radiatus* | MV403 | O | 120 | 22/04/2015 | 34,79 | -5,55 | Morocco |
| *Anacyclus* | *radiatus* | *radiatus* | MV409 | O | 17 | 22/04/2015 | 34,79 | -5,55 | Morocco |
| *Anacyclus* | *radiatus* | *radiatus* | MV413 | O | 152 | 22/04/2015 | 34,61 | -5,53 | Morocco |
| *Anacyclus* | *radiatus* | *radiatus* | MV54 | O | | 15/06/2014 | 31,50 | -9,70 | Morocco |
| *Anacyclus* | *radiatus* | *radiatus* | WPT44.1 | O | 134 | 19.06.2015 | 36,22 | -5,78 | Spain |
| *Anacyclus* | *valentinus* | | UPNA 11846 | UPNA | 164 | 24/05/2008 | 38,61 | -0,05 | Spain |
| *Anacyclus* | *valentinus* | | VIT 78628 | VIT | 207 | 2000 | 39,75 | -0,37 | Spain |
| *Anacyclus* | *valentinus* | | VIT 83639 | VIT | 188 | 2005 | 37,28 | -1,75 | Spain |
| *Anacyclus* | *valentinus* | | VIT 88094 | VIT | 159 | 2011 | 38,38 | -0,51 | Spain |
| *Anacyclus* | *valentinus* | | VIT 91146 | VIT | 212 | 1994 | 43,33 | -2,99 | Spain |
| *Matricaria* | *aurea* | | MV109 | O | 151 | 04/04/2015 | 31,01 | -7,23 | Morocco |
| *Matricharia* | *chamomilla* | | MV262 | O | 80 | 12/04/2015 | 31,95 | -8,24 | Morocco |
| *Otanthus* | *maritimus* | | MNHN-P-P04305585 | P | 41 | 2000 | 42,00 | 9,45 | Morocco |

#### Table S2.

| Collection ID | ID during sequencing | Date collected | GPS coodinates | | City | Country | Type of shop | Molecular identification | | |
|---|---|---|---|---|---|---|---|---|---|---|
| | | | ° N | ° E | | | | Genus | species | subsp. / var. |
| HAS425.2 | 7 | 2015 | 11,66 | 76,26 | Wayanad | India | Herbalist | *Anacyclus* | *pyrethrum* | *depressus* |
| MV503-1 | 8 | 01/06/2015 | 35,79 | -5,81 | Tanger | Morocco | Herbalist | *Anacyclus* | *pyrethrum* | *pyrethrum* |
| MV528.2 | 9 | 08/06/2015 | 33,60 | -7,62 | Casablanca | Morocco | Herbalist | *Anacyclus* | *pyrethrum* | *pyrethrum* |
| MV466A.2 | 13 | 22/04/2015 | 32,94 | -5,67 | Khenifra | Morocco | Herbalist | *Anacyclus* | *pyrethrum* | *depressus* |
| HAS438.2 | 14 | 2015 | 11,48 | 79,37 | Chidambaram | India | Herbalist | *Anacyclus* | *pyrethrum* | *depressus* |
| MV536-3.2 | 15 | 08/06/2015 | 33,60 | -7,62 | Casablanca | Morocco | Export company | *Anacyclus* | *pyrethrum* | *depressus* |
| HAS368.3 | 19 | 2015 | 14,47 | 75,92 | Davangere | India | Herbalist | *Anacyclus* | *pyrethrum* | *depressus* |
| MV501-3 | 20 | 01/06/2015 | 35,78 | -5,81 | Tanger | Morocco | Herbalist | *Anacyclus* | *pyrethrum* | *pyrethrum* |
| MV517-3 | 21 | 04/06/2015 | 34,03 | -6,84 | Rabat | Morocco | Traditional healer | *Anacyclus* | *pyrethrum* | *pyrethrum* |
| HAS368.1 | 24 | 2015 | 14,47 | 75,92 | Davangere | India | Herbalist | *Anacyclus* | *pyrethrum* | *depressus* |
| HAS418.1 | 26 | 2015 | 26,21 | 78,20 | Gwalior | India | Herbalist | *Anacyclus* | *pyrethrum* | *depressus* |
| MV536-6.2 | 27 | 08/06/2015 | 33,60 | -7,62 | Casablanca | Morocco | Export company | *Anacyclus* | *homogamos* | |
| MV466A.1 | 28 | 22/04/2015 | 32,94 | -5,67 | Khenifra | Morocco | Herbalist | *Anacyclus* | *pyrethrum* | *depressus* |
| HAS418.3 | 29 | 2015 | 26,21 | 78,20 | Gwalior | India | Herbalist | *Anacyclus* | *pyrethrum* | *depressus* |
| MV536-6.3 | 30 | 08/06/2015 | 33,60 | -7,62 | Casablanca | Morocco | Export company | *Anacyclus* | *homogamos* | |
| HAS421.1 | 32 | 2015 | 25,62 | 85,15 | Patna | India | Herbalist | *Anacyclus* | *pyrethrum* | *depressus* |
| MV539.1 | 33 | 01/04/2014 | 31,73 | -7,00 | Demnate | Morocco | Wholesaler | *Anacyclus* | *pyrethrum* | *depressus* |
| MV466B.3 | 34 | 22/04/2015 | 32,94 | -5,67 | Khenifra | Morocco | Herbalist | *Anacyclus* | *pyrethrum* | *depressus* |
| HAS421.2 | 35 | 2015 | 25,62 | 85,15 | Patna | India | Herbalist | *Anacyclus* | *pyrethrum* | *depressus* |
| MV539.4 | 36 | 01/04/2014 | 31,73 | -7,00 | Demnate | Morocco | Wholesaler | *Anacyclus* | *pyrethrum* | *depressus* |
| MV506-1.3 | 37 | 02/06/2015 | 33,89 | -5,57 | Meknes | Morocco | Traditional healer | *Anacyclus* | *pyrethrum* | *depressus* |
| MV501-1 | 38 | 01/06/2015 | 35,78 | -5,81 | Tanger | Morocco | Herbalist | *Anacyclus* | *pyrethrum* | *pyrethrum* |
| HAS408.2 | 43 | 2015 | 11,94 | 79,49 | Villupuram | India | Herbalist | *Anacyclus* | *pyrethrum* | *pyrethrum* |
| HAS433.1 | 52 | 2015 | 16,95 | 75,72 | peth vadgaon, kolhapur | India | Herbalist | *Anacyclus* | *pyrethrum* | *depressus* |
| MV510-1 | 53 | 03/06/2015 | 34,07 | -4,97 | Fès | Morocco | Traditional healer | *Anacyclus* | *pyrethrum* | *depressus* |
| MV528.3 | 54 | 08/06/2015 | 33,60 | -7,62 | Casablanca | Morocco | Herbalist | *Anacyclus* | *pyrethrum* | *pyrethrum* |
| MV466A.3 | 58 | 22/04/2015 | 32,94 | -5,67 | Khenifra | Morocco | Herbalist | *Anacyclus* | *pyrethrum* | *depressus* |
| MV536-3.3 | 60 | 08/06/2015 | 33,60 | -7,62 | Casablanca | Morocco | Export company | *Anacyclus* | *pyrethrum* | *depressus* |
| MV536-2.2 | 64 | 08/06/2015 | 33,60 | -7,62 | Casablanca | Morocco | Export company | *Anacyclus* | *pyrethrum* | *pyrethrum* |
| MV502-2 | 65 | 01/06/2015 | 35,79 | -5,81 | Tanger | Morocco | Herbalist | *Anacyclus* | *pyrethrum* | *pyrethrum* |
| MV520-3 | 66 | 05/06/2015 | 34,02 | -6,84 | Rabat | Morocco | Wholesaler, | *Anacyclus* | *pyrethrum* | *pyrethrum* |
| HAS398.1.2 | 68 | 2015 | 11,94 | 79,80 | Pondicherry | India | Herbalist | *Anacyclus* | *pyrethrum* | *depressus* |
| HAS374 | 69 | 2015 | 10,94 | 76,93 | Kovaipudur | India | Herbalist | *Anacyclus* | *pyrethrum* | *pyrethrum* |
| HAS434.3 | 76 | 2015 | 18,81 | 73,23 | Mumbai | India | Herbalist | *Anacyclus* | *pyrethrum* | *depressus* |
| MV517-2 | 77 | 04/06/2015 | 34,03 | -6,84 | Rabat | Morocco | Traditional healer | *Anacyclus* | *pyrethrum* | *pyrethrum* |
| MV534-2 | 78 | 08/06/2015 | 33,57 | -7,60 | Casablanca | Morocco | Export company | *Anacyclus* | *pyrethrum* | *pyrethrum* |
| MV300 | 79 | 18/04/2015 | 33,17 | -5,07 | Timadith | Morocco | Collector | *Anacyclus* | *pyrethrum* | *depressus* |
| MV466B.4 | 82 | 22/04/2015 | 32,94 | -5,67 | Khenifra | Morocco | Herbalist | *Anacyclus* | *pyrethrum* | *depressus* |
| MV536-4.1 | 84 | 08/06/2015 | 33,60 | -7,62 | Casablanca | Morocco | Export company | *Anacyclus* | *pyrethrum* | *depressus* |
| HAS403.1 | 88 | 2015 | 11,75 | 79,75 | Cuddalore | India | Herbalist | *Anacyclus* | *pyrethrum* | *depressus* |
| MV502-3 | 89 | 01/06/2015 | 34,79 | -5,81 | Tanger | Morocco | Herbalist | *Anacyclus* | *pyrethrum* | *pyrethrum* |
| HAS380.1 | 93 | 2015 | 10,26 | 78,89 | Trichy | India | Herbalist | *Anacyclus* | *pyrethrum* | *depressus* |
| HAS435.1 | 100 | 2015 | 22,53 | 88,34 | Kolkata | India | Herbalist | *Anacyclus* | *pyrethrum* | *depressus* |
| MV520-1 | 101 | 05/06/2015 | 34,02 | -6,84 | Rabat | Morocco | Wholesaler, | *Anacyclus* | *pyrethrum* | *pyrethrum* |
| MV535-2 | 102 | 08/06/2015 | 33,58 | -7,60 | Casablanca | Morocco | Export company | *Anacyclus* | *pyrethrum* | *pyrethrum* |
| MV308 | 103 | 18/04/2015 | 33,17 | -5,07 | Timadith | Morocco | Collector | *Anacyclus* | *pyrethrum* | *pyrethrum* |
| HAS398.3 | 106 | 2015 | 11,94 | 79,80 | Pondicherry | India | Herbalist | *Anacyclus* | *pyrethrum* | *depressus* |
| MV536-4.2 | 108 | 08/06/2015 | 33,60 | -7,62 | Casablanca | Morocco | Export company | *Anacyclus* | *pyrethrum* | *depressus* |
| HAS403.2 | 112 | 2015 | 11,75 | 79,75 | Cuddalore | India | Herbalist | *Anacyclus* | *pyrethrum* | *depressus* |
| MV503-3 | 113 | 01/06/2015 | 35,79 | -5,81 | Tanger | Morocco | Herbalist | *Anacyclus* | *pyrethrum* | *pyrethrum* |
| HAS435.2 | 124 | 2015 | 22,53 | 88,34 | Kolkata | India | Herbalist | *Anacyclus* | *pyrethrum* | *depressus* |
| MV526-1.1 | 125 | 05/06/2015 | 34,02 | -6,83 | Rabat | Morocco | Traditional healer | *Anacyclus* | *pyrethrum* | *pyrethrum* |
| MV536-1.2 | 126 | 08/06/2015 | 33,60 | -7,62 | Casablanca | Morocco | Export company | *Anacyclus* | *pyrethrum* | *pyrethrum* |
| MV309 | 127 | 18/04/2015 | 33,17 | -5,07 | Timadith | Morocco | Collector | *Anacyclus* | *pyrethrum* | *pyrethrum* |
| HAS425.1 | 130 | 2015 | 11,66 | 76,26 | Wayanad | India | Herbalist | *Anacyclus* | *pyrethrum* | *depressus* |
| MV514-2 | 131 | 03/06/2015 | 34,07 | -4,98 | Fès | Morocco | Traditional healer | *Anacyclus* | *pyrethrum* | *depressus* |
| MV540 | 132 | 01/04/2014 | 33,52 | -5,12 | Ifran | Morocco | Wholesaler | *Anacyclus* | *pyrethrum* | *depressus* |
| HAS430.2 | 136 | 2015 | 12,91 | 79,33 | Vellore | India | Herbalist | *Anacyclus* | *pyrethrum* | *depressus* |
| MV505-1 | 137 | 01/06/2015 | 35,78 | -5,81 | Tanger | Morocco | Herbalist | *Anacyclus* | *pyrethrum* | *pyrethrum* |
| MV526-1.3 | 138 | 05/06/2015 | 34,02 | -6,83 | Rabat | Morocco | Traditional healer | *Anacyclus* | *pyrethrum* | *pyrethrum* |
| HAS387.1 | 141 | 2015 | 8,49 | 76,95 | Thiruvanathapuram | India | Herbalist | *Anacyclus* | *pyrethrum* | *pyrethrum* |
| HAS438.1 | 148 | 2015 | 11,48 | 79,37 | Chidambaram | India | Herbalist | *Anacyclus* | *pyrethrum* | *depressus* |
| MV526-1.2 | 149 | 05/06/2015 | 34,02 | -6,83 | Rabat | Morocco | Traditional healer | *Anacyclus* | *pyrethrum* | *pyrethrum* |
| MV536-2.1 | 150 | 08/06/2015 | 33,60 | -7,62 | Casablanca | Morocco | Export company | *Anacyclus* | *pyrethrum* | *pyrethrum* |
| MV303 | 153 | 18/04/2015 | 33,17 | -5,07 | Timadith | Morocco | Collector | *Anacyclus* | *pyrethrum* | *depressus* |
| HAS425.4 | 154 | 2015 | 11,66 | 76,26 | Wayanad | India | Herbalist | *Anacyclus* | *pyrethrum* | *depressus* |
| MV514-3 | 155 | 03/06/2015 | 34,07 | -4,98 | Fès | Morocco | Traditional healer | *Anacyclus* | *pyrethrum* | *depressus* |
| MV541.1 | 156 | 05/09/2015 | 31,63 | -7,99 | Marrakech | Morocco | Herbalist | *Anacyclus* | *pyrethrum* | *depressus* |
| HAS433.2 | 160 | 2015 | 16,95 | 75,72 | peth vadgaon, kolhapur | India | Herbalist | *Anacyclus* | *pyrethrum* | *depressus* |
| MV506-2.2 | 161 | 02/06/2015 | 33,89 | -5,57 | Meknes | Morocco | Traditional healer | *Anacyclus* | *pyrethrum* | *pyrethrum* |
| MV534-3 | 162 | 08/06/2015 | 33,57 | -7,60 | Casablanca | Morocco | Export company | *Anacyclus* | *pyrethrum* | *pyrethrum* |
| MV502-1 | 165 | 01/06/2015 | 35,79 | -5,81 | Tanger | Morocco | Herbalist | *Anacyclus* | *pyrethrum* | *pyrethrum* |
| HAS439.1 | 172 | 2015 | 11,66 | 78,17 | Salem | India | Herbalist | *Anacyclus* | *pyrethrum* | *depressus* |
| MV526-2.2 | 173 | 05/06/2015 | 34,02 | -6,83 | Rabat | Morocco | Traditional healer | *Anacyclus* | *pyrethrum* | *pyrethrum* |
| MV536-2.3 | 174 | 08/06/2015 | 33,60 | -7,62 | Casablanca | Morocco | Export company | *Anacyclus* | *pyrethrum* | *pyrethrum* |
| MV302 | 177 | 18/04/2015 | 33,17 | -5,07 | Timadith | Morocco | Collector | *Anacyclus* | *pyrethrum* | *depressus* |
| HAS430.3 | 178 | 2015 | 12,91 | 79,33 | Vellore | India | Herbalist | *Anacyclus* | *pyrethrum* | *depressus* |
| MV541.3 | 180 | 05/09/2015 | 31,63 | -7,99 | Marrakech | Morocco | Herbalist | *Anacyclus* | *pyrethrum* | *depressus* |
| MV305 | 181 | 18/04/2015 | 33,17 | -5,07 | Timadith | Morocco | Collector | *Anacyclus* | *pyrethrum* | *pyrethrum* |
| HAS437.1 | 184 | 2015 | 12,95 | 74,87 | Mangalore | India | Herbalist | *Anacyclus* | *pyrethrum* | *depressus* |
| MV535-3 | 186 | 08/06/2015 | 33,58 | -7,60 | Casablanca | Morocco | Export company | *Anacyclus* | *pyrethrum* | *pyrethrum* |
| MV528.1 | 189 | 08/06/2015 | 33,60 | -7,62 | Casablanca | Morocco | Herbalist | *Anacyclus* | *pyrethrum* | *pyrethrum* |
| MV536-3.1 | 196 | 08/06/2015 | 33,60 | -7,62 | Casablanca | Morocco | Export company | *Anacyclus* | *pyrethrum* | *depressus* |
| MV542.2 | 198 | 05/09/2015 | 31,63 | -7,98 | Marrakech | Morocco | Herbalist | *Anacyclus* | *pyrethrum* | *pyrethrum* |
| HAS434.1 | 202 | 2015 | 18,81 | 73,23 | Mumbai | India | Herbalist | *Anacyclus* | *pyrethrum* | *depressus* |
| MV520-2 | 204 | 05/06/2015 | 34,02 | -6,84 | Rabat | Morocco | Wholesaler | *Anacyclus* | *pyrethrum* | *pyrethrum* |
| HAS437.3 | 208 | 2015 | 12,95 | 74,87 | Mangalore | India | Herbalist | *Anacyclus* | *pyrethrum* | *depressus* |
| MV536-1.3 | 210 | 08/06/2015 | 33,60 | -7,62 | Casablanca | Morocco | Export company | *Anacyclus* | *pyrethrum* | *pyrethrum* |
| MV536-5.2 | 216 | 08/06/2015 | 33,60 | -7,62 | Casablanca | Morocco | Export company | *Anacyclus* | *pyrethrum* | *depressus* |

| **Collection ID** | **ID during sequencing** | **Date collected** | **GPS coodinates** | | **City** | **Country** | **Type of shop** | **Putative identification (blastn)** | | |
|---|---|---|---|---|---|---|---|---|---|---|
| | | | **° N** | **° E** | | | | **Family** | **Genus** | **species** |
| MV466B.2 | 31 | 22/04/2015 | 32,94 | -5,67 | Khenifra | Morocco | Herbalist | Plantaginaceae | *Plantago* | spp. |
| MV600 | 39 | 09/08/2015 | 9,67 | 80,02 | Jaffna | Sri Lanka | Traditional healer | Plantaginaceae | *Plantago* | spp. |
| HAS372.2 | 40 | 2015 | 14,62 | 74,83 | Sirsi | India | Herbalist | Anthemidinae | | |
| MV601.1 | 42 | 15/08/2015 | 6,95 | 79,87 | colombo | Sri Lanka | Traditional healer | | | |
| MV507-2 | 44 | 02/06/2015 | 33,89 | -5,57 | Meknes | Morocco | Traditional healer | Plantaginaceae | *Plantago* | spp. |
| MV602 | 45 | 09/08/2015 | 9,67 | 80,02 | Jaffna | Sri Lanka | Traditional healer | Primulaceae | *Primula* | spp. |
| MV510-2 | 59 | 03/06/2015 | 34,07 | -4,97 | Fès | Morocco | Traditional healer | Anthemidinae | | |
| MV510-3 | 83 | 03/06/2015 | 34,07 | -4,97 | Fès | Morocco | Traditional healer | Anthemidinae | | |
| MV521-1 | 90 | 05/06/2015 | 34,03 | -6,84 | Rabat | Morocco | Traditional healer | Plantaginaceae | *Plantago* | spp. |
| MV467.2 | 92 | 22/04/2015 | 32,94 | -5,67 | Khenifra | Morocco | Herbalist | Plantaginaceae | *Plantago* | spp. |
| MV514-1 | 107 | 03/06/2015 | 34,07 | -4,98 | Fès | Morocco | Traditional healer | Anthemidinae | | |
| MV521-3 | 114 | 05/06/2015 | 34,03 | -6,84 | Rabat | Morocco | Traditional healer | Plumerieae | *Plumeria* | *spp.* |
| MV467.3 | 116 | 22/04/2015 | 32,94 | -5,67 | Khenifra | Morocco | Herbalist | Plantaginaceae | *Plantago* | spp. |
| HAS380.2 | 117 | 2015 | 10,26 | 78,89 | Trichy | India | Herbalist | | | |
| MV526-2.1 | 179 | 05/06/2015 | 34,02 | -6,83 | Rabat | Morocco | Traditional healer | Anthemidinae | | |
| MV507-1 | 185 | 02/06/2015 | 33,89 | -5,57 | Meknes | Morocco | Traditional healer | Plantaginaceae | *Plantago* | spp. |
| MV526-2.3 | 197 | 05/06/2015 | 34,02 | -6,83 | Rabat | Morocco | Traditional healer | | | |
| MV530 | 203 | 08/06/2015 | 33,60 | -7,62 | Casablanca | Morocco | Export company | Anthemidinae | | |
| MV511-1 | 209 | 03/06/2015 | 34,01 | -4,97 | Fès | Morocco | Traditional healer | Plantaginaceae | *Plantago* | spp. |
| MV504-3 | 213 | 01/06/2015 | 34,79 | -5,81 | Tanger | Morocco | Herbalist | Anthemidinae | | |
| HAS413.1 | 215 | 2015 | 11,77 | 79,55 | Panruti | India | Herbalist | Plantaginaceae | *Plantago* | spp. |

**Table S3.**

| Genus | Species | Collection ID - | Reference article |
|---|---|---|---|
| *Achillea* | *absinthoides* | AY603213.1 | 1 |
| *Achillea* | *acuminata* | AY603243.1 | 1 |
| *Achillea* | *ageratum* | AY603214.1 | 1 |
| *Achillea* | *alpina* | AY603206.1 | 1 |
| *Achillea* | *distans* | AY603188.1 | 1 |
| *Achillea* | *asiatica* | AY603208.1 | 1 |
| *Achillea* | *collina* | AY603198.1 | 1 |
| *Achillea* | *pratensis* | AY603199.1 | 1 |
| *Achillea* | *asplenifolia* | AY603201.1 | 1 |
| *Achillea* | *atrata* | AY603227.1 | 1 |
| *Achillea* | *ceretanica* | AY603203.1 | 1 |
| *Achillea* | *clavennae* | AY603229.1 | 1 |
| *Achillea* | *clusiana* | AY603228.1 | 1 |
| *Achillea* | *euxina* | AY603202.1 | 1 |
| *Achillea* | *fraasii* | AY603232.1 | 1 |
| *Achillea* | *holosericea* | AY603222.1 | 1 |
| *Achillea* | *lanulosa* | AY603205.1 | 1 |
| *Achillea* | *monticola* | AY603210.1 | 1 |
| *Achillea* | *oxyloba* | AY603238.1 | 1 |
| *Achillea* | *pannonica* | AY603184.1 | 1 |
| *Achillea* | *pyrenaica* | AY603247.1 | 1 |
| *Achillea* | *roseoalba* | AY603200.1 | 1 |
| *Achillea* | *salicifolia* | AY603244.1 | 1 |
| *Achillea* | *styriaca* | AY603190.1 | 1 |
| *Achillea* | *virescens* | AY603211.1 | 1 |
| *Achillea* | *wilsoniana* | AY603207.1 | 1 |
| *Anacyclus* | *clavatus* | AY603258.1 | 1 |
| *Anacyclus* | *monanthos* | KT954174.1 | 2 |
| *Anacyclus* | *pyrethrum* | KM887358.1 | 3 |
| *Anacyclus* | *pyrethrum* | KM887396.1 | 3 |
| *Anacyclus* | *pyrethrum* | KM887397.1 | 3 |
| *Anacyclus* | *pyrethrum* | KM887400.1 | 3 |
| *Heliocauta* | *atlantica* | AJ748782.1 | 4 |
| *Matricaria* | *aurea* | KT954177.1 | 2 |
| *Otanthus* | *maritimus* | AY603257.1 | 1 |
| *Tanacetum* | *argenteum* | AB683265.1 | 5 |
| *Tanacetum* | *aucherianum* | AB683268.1 | 5 |
| *Tanacetum* | *bachtiaricum* | AB683269.1 | 5 |
| *Tanacetum* | *balsamita* | AB683270.1 | 5 |
| *Tanacetum* | *bamianicum* | AB683271.1 | 5 |
| *Tanacetum* | *budjnurdense* | AB608330.1 | 5 |
| *Tanacetum* | *canescens* | AB608331.1 | 5 |
| *Tanacetum* | *chiliophyllum* | AB608332.1 | 5 |
| *Tanacetum* | *demetrii* | AB683278.1 | 5 |
| *Tanacetum* | *dumosum* | AB683281.1 | 5 |
| *Tanacetum* | *elbursense* | AB683282.1 | 5 |
| *Tanacetum* | *germanicopolitanum* | AB683285.1 | 5 |
| *Tanacetum* | *griffithii* | AB683286.1 | 5 |
| *Tanacetum* | *haradjanii* | AB683287.1 | 5 |
| *Tanacetum* | *haussknechtii* | AB683288.1 | 5 |
| *Tanacetum* | *hololeucum* | AB683289.1 | 5 |
| *Tanacetum* | *joharchii* | AB523746.1 | 5 |
| *Tripleurospermum* | *sp.* | KR150169.1 | 6 |

**Table S4.**

| | Target enrichment | | Genome skimming | |
|---|---|---|---|---|
| | Before Trimming | After trimming | Before Trimming | After trimming |
| Upper whisker | 7.20 | 6.30 | 6.80 | 6.20 |
| 3rd quartile | 4.20 | 3.70 | 4.10 | 3.60 |
| Median | 3.10 | 2.60 | 2.90 | 2.50 |
| 1st quartile | 2.20 | 1.90 | 2.20 | 1.80 |
| Lower whisker | 0.20 | 0.20 | 0.40 | 0.10 |
| Nr. of data points | 182 | 182 | 182 | 182 |
| Total reads | 626.5 | 542.6 | 587.2 | 507.2 |
| Average | 3.46 | 2.99 | 3.24 | 2.80 |

**Table S5.**

| | Coverage | | Alignment | |
|---|---|---|---|---|
| | Reference and trade samples | | Reference and trade samples | |
| | Target capture | Genome skimming | length (bp) | missing data (%) |
| Nuclear markers | 303 | 12 | 289.236 | 1.2% |
| Plastome | - | 20 | 110.003 | 2.1% |
| nrDNA | - | 131 | 633 | 2.7% |
| *matk* | - | 20 | 1523 | 0.0% |
| *rbcl* | - | | 1438 | 0.0% |
| *trnH-psbA* | - | | 500 | 0.3% |
| *trnL* | - | | 947 | 4.6% |

**Table S6.**

| | Adulteration | A. pyrethrum var. pyrethrum | A. pyrethrum var. depressus | A. homogamos |
|---|---|---|---|---|
| Collectors | 0% | 63% | 38% | 0% |
| Export company | 0% | 50% | 38% | 13% |
| Herborist - India | 22% | 14% | 63% | 0% |
| Herborist - Morocco | 32% | 46% | 22% | 0% |
| Traditional healer - Morocco | 57% | 22% | 21% | 0% |
| Wholesaler | 2% | 31% | 67% | 0% |

### Table S7.

| | N.A. | | Genus | | Species | | Population | |
|---|---|---|---|---|---|---|---|---|
| | percentage | # indiv. | percentage | # indiv. | percentage | # indiv. | percentage | # indiv. |
| *rbcl* | 100% | 110 | 0% | 0 | 0% | 0 | 0% | 0 |
| *trnH-psbA* | 100% | 110 | 0% | 0 | 0% | 0 | 0% | 0 |
| *matk* | 45% | 49 | 55% | 61 | 0% | 0 | 0% | 0 |
| *trnL* | 54% | 59 | 46% | 51 | 0% | 0 | 0% | 0 |
| ITS | 7% | 8 | 93% | 102 | 0% | 0 | 0% | 0 |
| Plastome | 44% | 48 | 56% | 62 | 6% | 7 | 0% | 0 |
| Nuclear markers | 9% | 10 | 81% | 89 | 81% | 89 | 55% | 60 |
